# Adulthood depletion of Integrator extends lifespan and healthspan via defective pre-mRNA processing

**DOI:** 10.64898/2026.04.18.719358

**Authors:** Luke Slade, Krzysztof Kuś, Yuanhang Jiang, Juan Cabello, Milos Filipovic, Peter Sarkies, Lidia Vasiljeva

## Abstract

Identifying strategies to mitigate age-related physiological decline remains a central challenge. During ageing, the transcriptome undergoes extensive remodelling, but how this affects organismal health and lifespan is not well understood. The Integrator complex plays a central role in RNA polymerase II transcription and RNA 3′ end processing. Surprisingly, we find that depletion of most Integrator subunits specifically in adults extends lifespan and healthspan in the nematode *C. elegans*. We show that loss of the catalytic subunit INTS-11 disrupts 3′ end formation of small nuclear and spliced leader RNAs, impairing trans-splicing and promoting outron retention in a subset of transcripts enriched for spliceosomal and nucleocytoplasmic transport genes. These RNA-processing defects lead to altered levels of endogenous siRNAs, which are required for the longevity and healthspan benefits of INTS-11 depletion. In parallel, outron retention disrupts nuclear-encoded mitochondrial gene expression and protein production, inducing mitochondrial dysfunction and promoting lifespan extension. We also demonstrate that loss of INTS-11 perturbs transcription elongation at genes where Integrator is present at promoters, and that upregulation of enhancer elements within intragenic regions can affect the expression and isoform usage of nearby genes. Together, our findings identify Integrator as a key upstream regulator of non-coding RNA transcription, which in turn impacts protein-coding gene expression and mitochondrial function to shape the ageing process.

## Introduction

Ageing and its associated co-morbidities place increasing strain on healthcare systems and public finances. In England alone, the population aged ≥65 years is projected to rise by 3.3 million within 20 years and by 6.5 million within 40 years [1]. Identifying interventions that attenuate physiological decline (i.e., healthspan) rather than lifespan alone is therefore a pressing socio-economic priority.

At the molecular level, many hallmarks of ageing converge on altered gene expression programs [2, 3]. In metazoans, RNA polymerase II (Pol II) transcribes both protein-coding and non-coding genes and coordinates co-transcriptional RNA processing, including 5′ capping, pre-mRNA splicing, and 3′ end formation [4], events tightly coupled to transcription [5]. Importantly, transcriptional and RNA-processing defects have been linked to cellular and physiological deterioration across species [6, 7]. Yet, whilst essential during developmental periods, how specific Pol II-associated regulatory complexes shape organismal ageing, particularly in adulthood, remains poorly defined.

One such complex is Integrator, a conserved, multi-subunit complex that is a central mediator of inducible transcriptional programs [8–15]. Integrator was originally implicated in small nuclear RNA (snRNA) 3′ end processing [16], but later studies showed that it also regulates transcription at protein-coding genes by attenuating promoter-proximally paused Pol II and limiting progression of elongation-incompetent complexes into gene bodies [10, 17, 18]. Catalytic activity of the endonuclease subunit INTS11 is central to Integrator’s function, mediating cleavage of the nascent RNAs 3′ end [9, 16, 19] and promoting transcription termination [20, 21]. Additionally, the Integrator-associated PP2A phosphatase promotes transcriptional attenuation by dephosphorylating Pol II and elongation factors. This opposes CDK9 (P-TEFb)-mediated phosphorylation of Spt5, Pol II, and the negative elongation factor (NELF) known to promote pause release [22–25].

In addition to snRNA, Integrator is essential for 3’ end formation of many other non-coding transcripts including telomerase RNA (TERC), enhancer RNA (eRNA), NEAT1 RNA and small RNAs including viral microRNAs and piRNA [26–33]. Despite increasing interest and advances on the structural, biochemical and genomic aspects of Integrator [25, 34–36], little is known about the role of this essential complex in adult physiology and ageing [37, 38]. In humans, Integrator dysfunction has been linked to neurodevelopmental disorders and cancer [39–41]. Conversely, elevated expression of specific subunits correlates with poor prognosis in several malignancies, where silencing reduces pathological markers [42–44]. These observations suggest that Integrator may have previously unrecognised post-developmental roles in regulating organismal healthspan.

To investigate post-developmental functions in a whole-organism context, tractable *in vivo* systems are essential. In the pursuit of accelerating healthspan research, the simple model organism *Caenorhabditis elegans* has played a central role in uncovering seminal conserved mechanisms of age-associated decline [45–47]. With its short ∼3-week lifespan, ∼85% genomic conservation with humans, translucent body and powerful genetic toolkit, *C. elegans* enables precise genotype-to-phenotype mapping across the life course. Indeed, studies in *C. elegans* have already contributed to our understanding of Integrator function, revealing developmentally essential subunits [19] and a catalytic requirement for piRNA biogenesis [32, 48]. What is lacking, however, is how Integrator contributes to the ageing process during adulthood, where *C. elegans* provides a candidate model to address this.

Here, we depleted Integrator subunits in adult worms to define their contributions to ageing. Strikingly, we show that silencing of the majority of Integrator subunits, notably including the catalytically active subunit extends lifespan. We demonstrate this effect arises from widespread transcriptional dysfunction upon loss of Integrator. In particular, impaired biogenesis of splice-leader RNAs (SL RNAs) interferes with trans-splicing, resulting in the production of small RNAs that target misprocessed genes. Crucially, these effects converge on reduced mitochondrial protein levels, extending lifespan through impaired mitochondrial function. Our work identifies a previously unknown role for Integrator in the control of lifespan and healthspan.

## Results

### Loss of Integrator impacts organismal lifespan and healthspan

The mammalian Integrator complex comprises 15 subunits organised into structurally and functionally distinct sub-complexes (Fig. 1A). *C. elegans* has retained at least 13 subunits with conserved molecular functions. Although many Integrator subunits are essential for development [19], depletion of essential genes can also exert different phenotypes during ageing [49–53] suggesting that their function in a context of adult physiology may differ (i.e., antagonistic pleiotropic (AP) gene function [54])). To assess adult-specific functions of Integrator, we depleted individual subunits by RNAi from the L4 stage onwards. Knockdown of *ints-1/3/7* shortened lifespan and/or healthspan, RNAi against *ints-9* had no effect, whereas depletion of 8 of 12 subunits extended lifespan (*ints-2/4/5/6/8/11/12/13*; Fig. 1B-C and Fig. S1A). We decided to focus on the core catalytic subunit *ints-11* to investigate mechanisms of its role in longevity.

**Figure 1.**
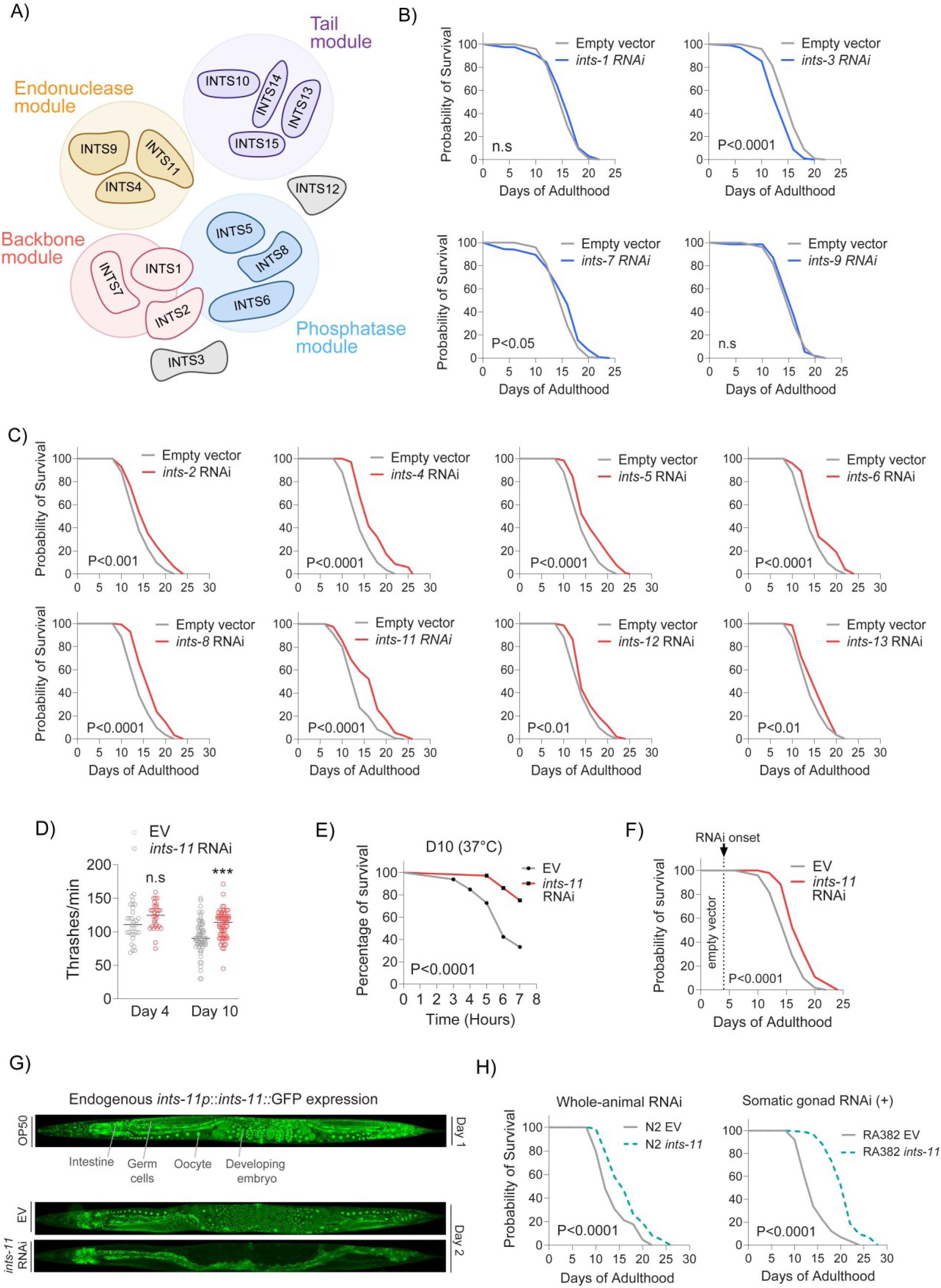
Post-developmental depletion of Integrator extends lifespan preferentially within the somatic gonad. **A)** Model of Integrator complex highlighting proposed sub-complex formations. **B)** Integrator subunits whereby targeted RNAi (starting from young-adulthood) either shortens or has no effect on animal lifespan. **C)** Integrator subunits whereby lifespan is significantly extended under targeted RNAi (from young-adulthood). **D)** Neuromuscular thrashing tests 4- and 10-days post *ints-11* RNAi onset. **E)** Thermotolerance tests 10-days post *ints-11* RNAi induction. **F)** Lifespan comparing empty vector controls against *ints-11* RNAi starting exclusively from day-4 of adulthood (post-reproductive period). **G)** Fluorescence microscopy of an endogenously tagged INTS-11::GFP strain in day-1 adults (top). Comparison of INTS-11::GFP signal localisation after animals were fed either empty vector bacteria for 2-days or dsRNA targeting *ints-11* (bottom). Note organism-wide depletion of INTS-11 from dsRNA feeding targeting *ints-11*. **H)** Lifespan is extended further under tissue-specific silencing of *ints-11* within the somatic gonad compared to whole-animal knockdown. All lifespans are from 2 independent repeats, with ∼200 animals per condition. Thrashing assays are from 2 independent repeats, with ∼ 50 animals per condition. INTS-11 images are representative of ∼20 individual animals per condition.

*ints-11* depletion also improved healthspan, enhancing neuromuscular activity (Fig. 1D) and thermotolerance in aged animals (Fig. 1E). As reduced reproduction is often linked to lifespan extension [55–58], we tested whether altered fecundity contributed to observed phenotypic effects. *ints-11* depletion from L4 reduced viable F1 progeny on days 1-2 of adulthood (Fig. S1B). To directly test a possible reproductive trade-off, we used the temperature-sensitive *glp-1*(*e2141*) mutant, in which shifting animals from 20°C to 25°C during development ablates the germline and renders adults sterile. *ints-11* RNAi continued to extend lifespan in sterile *glp-1*(*e2141*) animals (Fig. S1C). Moreover, initiating *ints-11* RNAi from day-4 of adulthood (after reproduction has largely ceased) still significantly extended lifespan (Fig. 1F). Together, these data indicate that reducing post-developmental levels of *ints-11* promotes longevity independently of its effects on reproduction.

### *ints-11* depletion acts primarily within the somatic gonad to attenuate ageing decline

To determine the tissues responsible for longevity, we generated an endogenous INTS-11::GFP reporter using CRISPR-Cas9 to monitor tissue distribution and RNAi-dependent loss. Fluorescence imaging revealed broad INTS11 expression, largely nuclear across tissues, with the strongest signal in developing embryos, oocytes, and germ cells (Fig. 1G). This signal disappeared completely after two days of RNAi exposure, confirming effective depletion of INTS-11 throughout the entire animal (Fig 1G). To probe if certain tissues were responsible for longevity phenotypes, we performed tissue-specific RNAi screening. Depletion of *ints-11* in the germline or hypodermis did not significantly alter survival (Fig. S1D), and body-wall muscle depletion shortened lifespan (Fig. S1D). Silencing within dopaminergic, glutaminergic and cholinergic neuronal strains revealed no changes (Fig. S1D), whereas intestinal depletion produced moderate lifespan extension (Fig. S1D). The most pronounced extension of lifespan was observed when *ints-11* was depleted in the somatic gonad (Fig. 1H), corroborated by neuromuscular activity improvements (Fig. S1E). Thus, the degree of lifespan extension observed in wild-type animals likely reflects divergent tissue-specific requirements for *ints-11*, with the somatic gonad contributing most strongly to the longevity phenotype.

### Depletion of *ints-11* elicits protein-coding transcriptome remodelling

We hypothesised that alterations in gene regulation might be responsible for *ints-11* depletion-induced longevity. To test this, we first performed INTS-11 chromatin immunoprecipitation followed by sequencing (ChIP–seq) in day-1 adult *C. elegans* using a 3xFLAG-tagged strain to determine possible sites where INTS-11 is recruited genome-wide.

We identified 7,276 high-confidence peaks and annotated them using a recently published comprehensive map of genome genic (non-coding and protein-coding) regulatory regions [59]. Of these, 649 mapped to protein-coding promoters, 447 to putative enhancers, 24 to non-coding RNA loci, 11 to pseudogene promoters and 33 to unassigned promoter regions, with a further 176 overlapping previously undefined regulatory elements (Fig. S2A). To identify further potential loci bound by Integrator, we aligned all remaining peaks to the canonical WBcel235 genome annotation. This revealed substantial intragenic occupancy, including 2,183 peaks within exons, 1,169 within introns, 498 at transcription start sites (TSS), 765 at transcription end sites of protein-coding genes (TES) and 1,403 in intergenic regions (Fig. 2A). Peaks with the highest read coverage were enriched at TSS and TES of protein-coding genes (Fig. 2B). Together, these data indicate that, unlike the predominantly promoter-proximal binding enrichment observed in mammals, INTS-11 in *C. elegans* exhibits extensive intragenic binding, suggesting divergence in genomic targeting across species.

**Figure 2.**
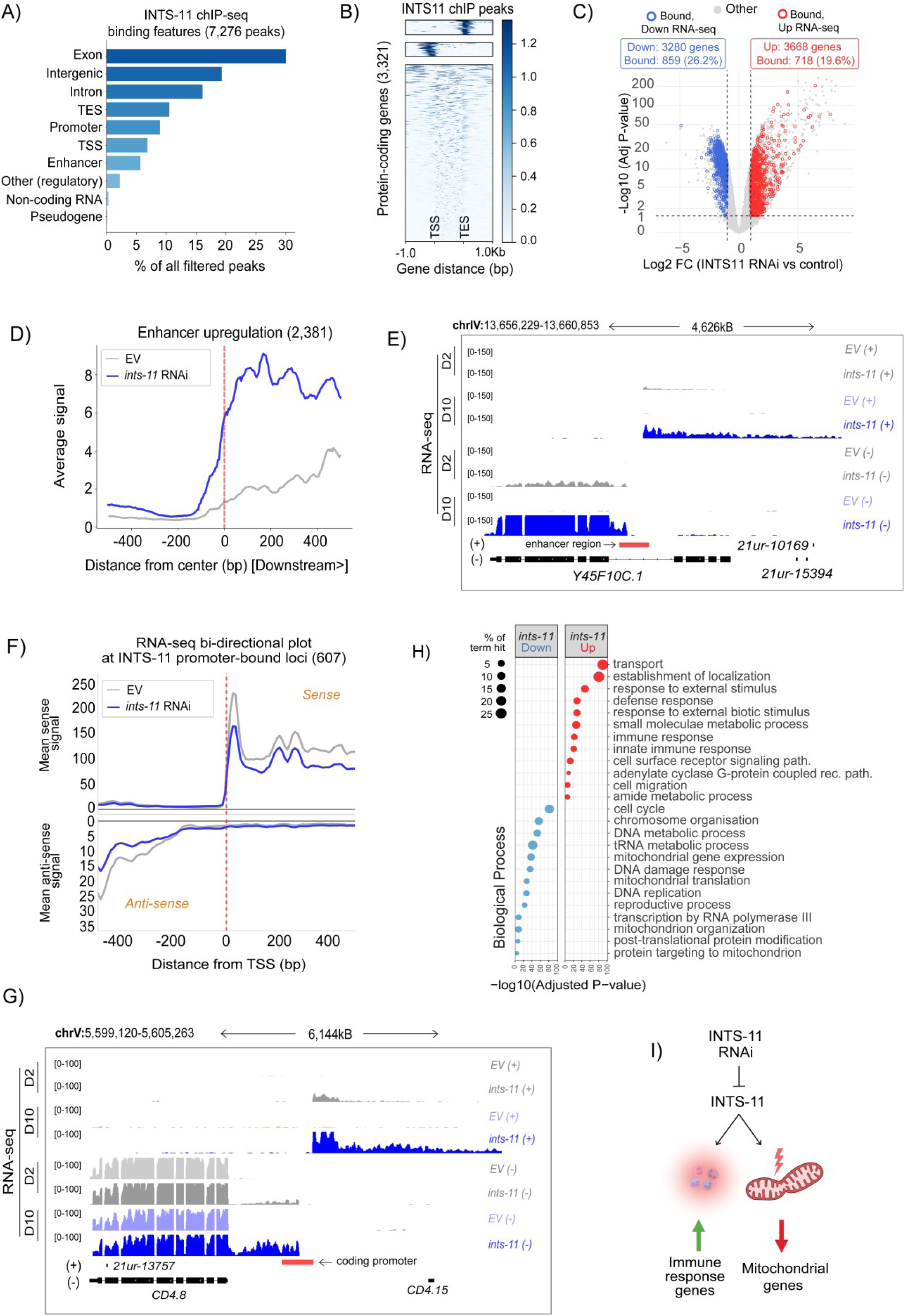
The INTS-11 binding landscape and transcriptomic re-modelling from depletion of *ints-11* in aged animals. **A)** Summary bar plot of INTS11-bound genomic features in day-1 adults (7,276 genes total). **B)** Heatmap of INTS11 ChIP-seq peaks across protein-coding genes (2,933), filtered for divergent or tandem genes (300bp window) and clustered into three distinct genomic features (TSS, TES and intragenic). **C)** Volcano plot representing differential gene expression from ribodepleted RNA-seq of day-10 aged animals. Genes identified as bound from INTS-11 ChIP-seq are shown in blue (bound + downregulated RNA) and red (bound + upregulated RNA). **D)** Metagene illustrating upregulated gene expression at enhancer regions from *ints-11* silencing in day-10 adults. **E)** Genome coverage tracks of a representative enhancer region bound by INTS11, exhibiting upregulated gene expression from *ints-11* RNAi. Note upregulation is present in animals fed *ints-11* RNAi for only 2-days, becoming more severe in day-10 aged animals. **F)** Metagene plot of RNA-seq coverage +/- 400bp around INTS-11-bound protein-coding promoter regions identified in ChIP-seq datasets. **G)** Genome coverage tracks of a representative bidirectional promoter upregulation event. **H)** Gene ontology (biological process) enrichment plot of all genes downregulated (3,280) and upregulated (3,668) in *ints-11* RNAi. **I)** Summary schematic representing the primary enriched up/ downregulated gene expression pathways from *ints-11* RNAi in aged animals.

To determine how INTS11 occupancy related to transcriptional changes underlying longevity, we performed high-throughput sequencing of ribodepleted RNA from day-10 adults under *ints-11* RNAi, identifying 6,948 differentially expressed genes (3,280 downregulated and 3,668 upregulated; Fig. 2C and supplemental table 1). We then compared expression changes across coding sequence regions between INTS11-bound and unbound genes, and found INTS11 occupancy did not correlate with genome-wide RNA expression changes following depletion (Fig. S2B). We also did not observe differences in metagene profiles between bound and unbound loci within genes that were either upregulated or downregulated at the RNA level (Fig. S2C). These results suggest that the remodelling of the protein-coding transcriptome upon *ints-11* knockdown likely reflects secondary transcriptional effects rather than direct binding-dependent regulation.

### Enhancer and bidirectional promoter transcription is associated with genes outside of longevity-enriched pathways

Integrator has been implicated to function at genic promoters to suppress antisense transcription and control promoter-proximal pausing in mammalian cells [18, 23, 60]. Spt5 and NELF are proposed to mediate Integrator function in promoter proximal pause regulation [34, 61]. Interestingly, NELF is not evolutionary conserved in *C. elegans* [62], making it a great system to understand mechanisms of Integrator function. We therefore assessed transcriptional changes at enhancer and promoter regions to assess whether Integrator function, despite the lack of NELF, is conserved at promoter and enhancer regions. We found depletion of *ints-11* led to enhancer RNA upregulation at 2,381 loci (Fig. 2D and supplemental table 2), an effect that was observed irrespective of their basal transcriptional activity (Fig. S2D). Of these, 1,753 upregulated enhancers were located within intragenic regions: 318 and 287 of these intragenic enhancer-associated genes were found to be up- and downregulated, respectively, in *ints-11* depleted animals (supplemental table 2). Strikingly, we demonstrate that enhancer activation in intragenic regions shows upregulation of what appears as an alternative mRNA isoform at a subset of loci (representative example in Fig. 2E). Consistently, intragenic enhancers were shown to regulate gene expression on mammalian cells by acting as alternative promoters to generate spliced, multi-exonic, polyadenylated transcripts with altered coding potential [63]. Additionally, activation of intragenic enhancers can, in some cases, interfere with host gene transcription [64]. Our data demonstrate that Integrator can regulate gene expression by modulating expression of enhancers.

Loss of Integrator in mammals is known to produce promoter-upstream transcripts (PROMPTs), unstable transcripts resulting from bi-directional promoters [21]. Importantly, previous work has shown that promoter-associated antisense transcription is widespread in *C. elegans* [65]. To investigate whether Integrator functions in the suppression of promoter-driven antisense transcription, we assessed gene expression coverage at the promoters of genes bound by INTS-11. We found no strong evidence for upregulated antisense transcription at these loci in *ints-11* depleted samples (Fig. 2F). When performing the same analysis genome-wide, we identified 465 genes with upregulated antisense transcription, consistent with a conserved role for Integrator in the suppression of PROMPTs (Fig. 2G and S2E and supplemental table 1). To note, the number of these events is likely limited by the fact PROMPTs are unstable in nature, and are rapidly degraded by the RNA exosome machinery [66, 67] and thus capable of evading detection by total RNA-seq. Furthermore, promoter occupancy of Integrator correlates with reduced transcription elongation (Fig. 2F), consistent with studies in mammalian systems showing that Integrator regulates elongation-incompetent Pol II complexes and that its loss leads to widespread elongation defects [27]. This contrasts with findings in *Drosophila*, where Integrator acts at promoters to attenuate gene expression via premature transcription termination, resulting in increased transcription upon Integrator depletion [8, 18]. We suggest, as has been reported in mammalian systems, Integrator might be less involved in premature transcription-termination, but rather as a quality control checkpoint for competent Pol II transcription into gene bodies in *C. elegans* [27].

To understand how genes associated with enhancer and bidirectional promoter-upregulation might underpin longevity phenotypes, we performed gene ontology enrichment analysis on all differentially expressed genes identified in RNA-seq analysis (3,280 downregulated and 3,668 upregulated). We observed a striking enrichment for mitochondria-related process amongst downregulated genes such as mitochondrial gene expression, translation, organisation and targeting to mitochondria (Fig. 2H), with many genes belonging to immune response processes amongst upregulated genes (Fig. 2H-2I and S2F). Importantly, genes proximal to upregulated enhancer and bidirectional promoter regions were enriched for developmental processes including cell cycle regulation, G1/S transition, and embryo development (Fig. S2G). Thus, genes in close proximity to bidirectional promoter and enhancer activity represent a functionally distinct subset, separate from the broader biological pathways identified across all differentially expressed transcripts (Fig. 2H).

### *ints-11* depletion induces defective transcription-termination at non-coding loci

A key function of Integrator in both mammals and *C. elegans* is termination of small non-coding RNA transcription. Consistent with this, *ints-11* RNAi induced pronounced termination defects at snRNA loci (Fig. 3A–B) and, to a lesser extent, snoRNA loci (Fig. S3A–S3D). Readthrough extended substantially beyond annotated TES: among the 56 snRNA genes with the strongest defects, ∼50% exhibited Pol II readthrough up to 5 kb downstream, with ∼20% extending as far as 15 kb (Fig. 3C) spanning regions that can include several genes (Fig. 3B). While read-through transcription into neighbouring genes is associated with increased signal, these extended reads have been shown to produce long chimeric aberrant transcripts that are subjected to rapid degradation by nuclear surveillance complexes and represses transcription from the down-stream promoters due to transcription interference [67–69].

**Figure 3.**
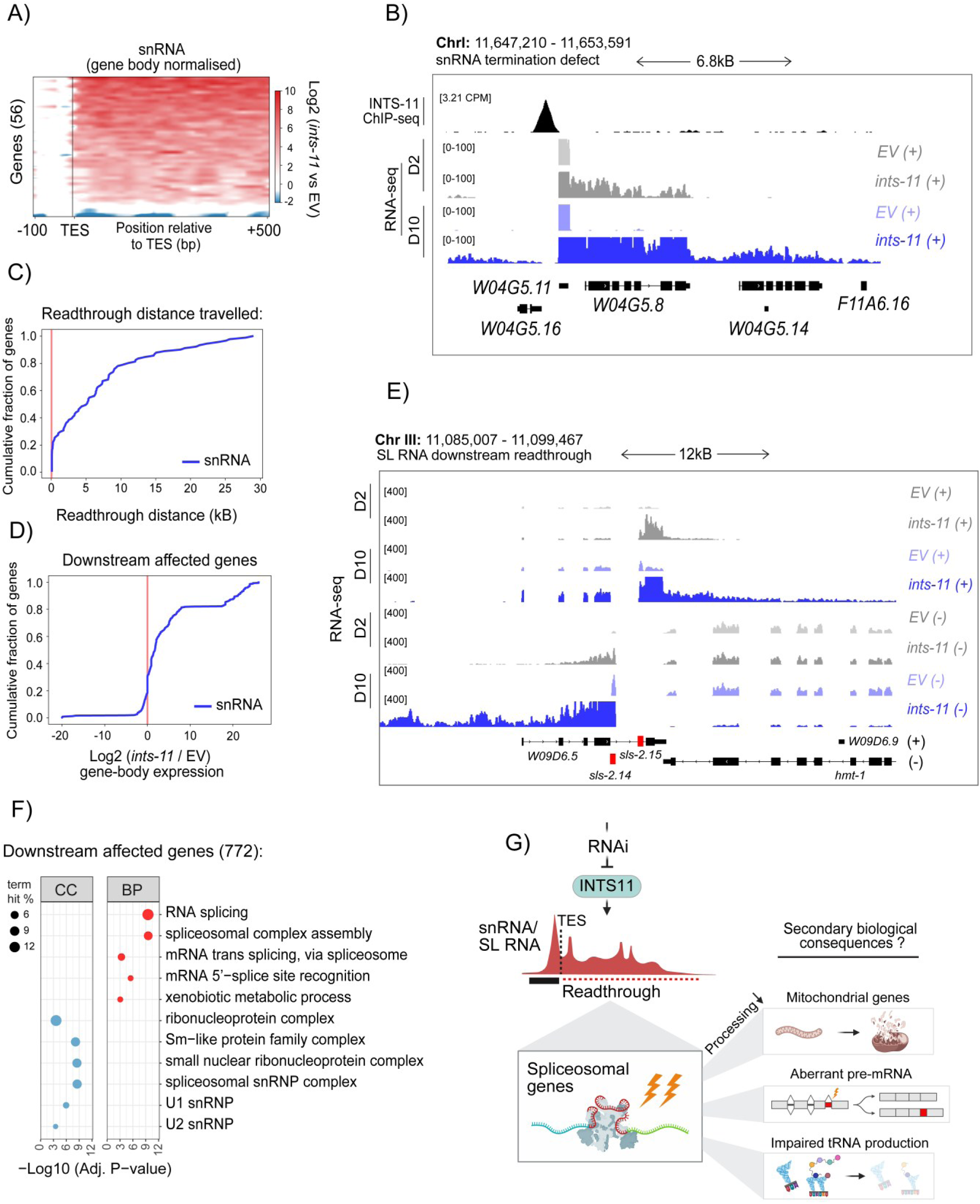
Spliceosomal genes are disproportionately affected by Pol II readthrough from defective snRNA transcription termination in *ints-11*-depleted animals. **A)** Heatmap of gene-body normalised read coverage downstream (500bp window) of snRNA transcription termination sites. A region of - 100bp is shown upstream of the termination site to show normalised gene-body signal. **B)** Representative genome coverage tracks of an INTS11-bound U2 snRNA (*W04G5.11*) showing extensive readthrough in animals fed *ints-11* RNAi. Note, termination defects become more severe in day-10 *vs* day-2 silenced animals. **C)** Cumulative distribution plot of total genomic distance travelled by each snRNA exhibiting defective transcription termination. **D)** Cumulative distribution plot of Log_2_ (*ints-11* RNAi *vs* EV) gene abundancies in genes residing within downstream snRNA readthrough windows. **E)** Genome coverage tracks of two SL RNAs (marked red), highlighting the consequences of transcription termination defects on neighbouring genes across both DNA strands. **F)** Gene ontology enrichment (biological process) for genes affected by downstream transcriptional readthrough. **G)** Proposed model in which *ints-11* depletion disrupts snRNA transcription termination, resulting in pol II readthrough into adjacent spliceosomal genes which initiates secondary transcriptional perturbations on distinct pathways.

Next, to determine the impact on neighbouring genes, we defined genomic intervals corresponding to each readthrough window and quantified expression of the downstream genes across both DNA strands. This revealed a significant increase in gene-body coverage of the downstream genes following *ints-11* depletion (Fig. 3D–E), reflective of non-productive readthrough transcription rather than bona fide promoter-driven expression. Interestingly, genes affected by readthrough were strongly enriched for spliceosomal functions, including trans-splicing, Sm-protein complexes and snRNP assembly (Fig 3F). Operon-like gene organisation is common in *C. elegans*, with ∼15–20% of genes arranged in operons and transcribed as polycistronic units that are resolved by trans-splicing [70, 71]. Genes within operons are typically closely spaced and often functionally related, with downstream genes receiving the SL2 spliced leader. These operons are enriched for housekeeping and RNA-processing functions, including splicing factors, consistent with the clustering of spliceosomal genes observed downstream of snRNA loci in our analysis. Together, these data indicate that defective snRNA termination drives transcriptional readthrough across loci encoding for factors involved in splicing, interfering with their expression (Fig. 3G).

### Defective trans-splicing leads to genome-wide outron retention linked to longevity

To assess how loss of *ints-11* affects pre-mRNA splicing, we applied rMATS-turbo to day-10 RNA-seq samples, a gold-standard bioinformatic pipeline for alternative-splicing analysis [72]. Changes were modest, with 526 significant events (adjusted P<0.05; Δ percentage spliced-in ± 10%) spanning alternative 3′ splice sites (290), 5′ splice sites (18), exon skipping (154), intron retention (43) and mutually exclusive exons (21; Fig. S4A-C), predominantly affecting developmental genes (Fig. S4D). However, beyond cis-splicing, *C. elegans* relies extensively on trans-splicing (∼85% of genes), in which an SL RNA donates its 5′ exon to pre-mRNAs, replacing the native 5′ UTR to promote transcript maturation and stability [70, 71, 73, 74]. Failure of trans-splicing results in retention of the native 5′ region, termed hereafter as the outron. Using total RNA-seq, we detected termination defects across the SL1 RNA cluster required for SL1-mediated trans-splicing (Fig. 4A). Consistent with impaired SL RNA transcription, we observed widespread outron retention affecting 2,631 genes (Fig. 4B and supplementary table 3). Outron-retaining genes were enriched for spliceosomal and nucleocytoplasmic transport pathways (Fig. 4C-E), suggesting a feed-forward disruption in which defective trans-splicing compromises expression of genes required for snRNP assembly, import and subsequent (trans)splicing reactions (Fig. 4F).

**Figure 4.**
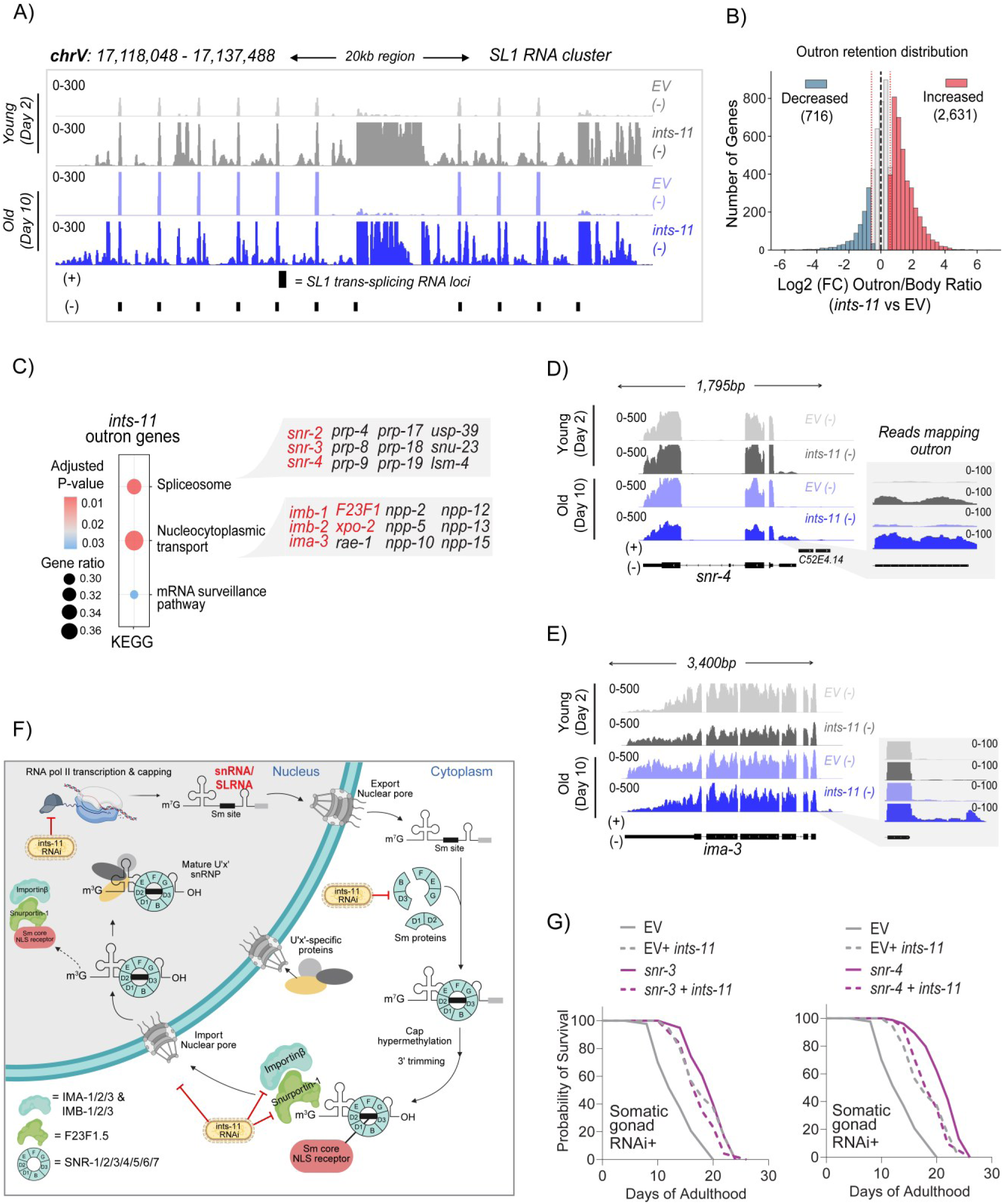
Loss of *ints-11* perturbs outron removal of spliceosomal and nuclear transport genes, promoting lifespan extension via defective trans-splicing. **A)** Genome coverage tracks of the SL1 RNA cluster required for trans-splicing, highlighting the defective transcription termination of SL1 RNAs in animals fed *ints-11* RNAi 2- and 10-days post adulthood. **B)** Distribution plot showing increased (3,889) number of genes with reads mapping to outron regions of pre-mRNAs in *ints-11* silenced animals. **C)** KEGG pathway enrichment of outron-retained genes in *ints-11* silenced animals, enriched for spliceosomal, nucleocytoplasmic transport and mRNA surveillance pathways. **D)** Representative genome coverage tracks of representative Sm protein (*snr-4*/ Sm D2) and nuclear importin (E; *ima-3*/ KPNA3) genes with significant outron retention. **F)** Pathway model of snRNP biogenesis steps affected by *ints-11* silencing. SL and snRNAs are transcribed in the nucleus, exported to the cytoplasm, assembled with Sm proteins, processed at their 5′ cap and 3′ end, and then re-imported to the nucleus as mature snRNPs to support pre-mRNA splicing. Upon *ints-11* depletion, snRNA/SL RNA transcription fails to terminate properly, and we detect retention of 5′ UTR outron sequences (a hallmark of impaired trans-splicing) in mRNAs encoding factors across the snRNP biogenesis pathway (e.g., Sm proteins and nuclear import components). We propose that termination defects reduce effective snRNP production, weaken splicing/trans-splicing and thereby amplify outron retention in snRNP-biogenesis genes in a feed-forward loop. **G)** Adulthood RNAi against either *snr-3* or *snr-4* within the *ints-11-*responsive tissue (somatic gonad) dramatically extends lifespan, which is not additive with combined *ints-11* silencing, implying genetic epistasis for *ints-11*-mediated longevity with defective trans-splicing.

If altered expression of SL RNAs were important for lifespan extension due to *ints-11* knockdown, then we would not expect to see defects in SL RNA production in knockdowns of Integrator subunits that do not extend lifespan. We therefore examined published RNA-seq data from *ints-3* RNAi knockdown [19], representing a subunit with compromised lifespan and healthspan phenotypes upon depletion in adulthood (see Fig. 1A and S1A). *ints-3* knockdown did not result in noticeable termination defects at SL RNA loci, suggesting that this effect could be specifically associated with lifespan-extending subunits (Fig. S4E-G).

To further test the contributions of small RNAs to *ints-11*-mediated lifespan extension, we targeted two components of the Sm-core gene family (*snr-3* and *snr-4*) involved in SL and snRNP biogenesis [75–77] which have previously been reported to block trans-splicing action [78]. We found that knockdown of both Sm genes within the *ints-11*-responsive tissue (somatic gonad) induced lifespan extension. Importantly, knockdown of Sm genes and *ints-11* together did not further extend lifespan (Fig. 4G). Together, these data support the hypothesis that *ints-11* knockdown extended lifespan through reduced expression of splicing components.

### Impaired mitochondrial and spliceosomal protein levels in *ints-11*-depleted animals

As trans-splicing defects are predicted to impair mRNA maturation and translation [73, 79], we examined the ageing proteome following *ints-11* depletion. We identified 845 differentially expressed proteins as predominantly downregulated in day-10 adults (768 decreased versus 77 increased; Fig. 5A and supplementary table 4). Importantly, outron-retaining transcripts were significantly associated with reduced protein abundance (Fig. S5A). Gene ontology analysis revealed enrichment of spliceosomal (U1 and U4 snRNP) pathways among downregulated proteins (Fig. 5B), consistent with RNA-seq and outron-retention datasets. Notably, abundancies of all Sm proteins required for SL and snRNP assembly (SNR-1/2/3/4/5/6/7) were reduced (Fig. S5B). Proteins involved in nucleocytoplasmic transport were also diminished, including snRNP biogenesis factors required for SL and snRNA nuclear re-import (IMB-3, IMA-2, IMA-3; Fig. S5B), earlier identified as affected by outron retention. Together, these data demonstrate that defective trans-splicing associates with reduced expression of proteins required for snRNP import and subsequent splicing reactions, reinforcing a positive-feedback loop that further disrupts outron removal and protein production.

**Figure 5.**
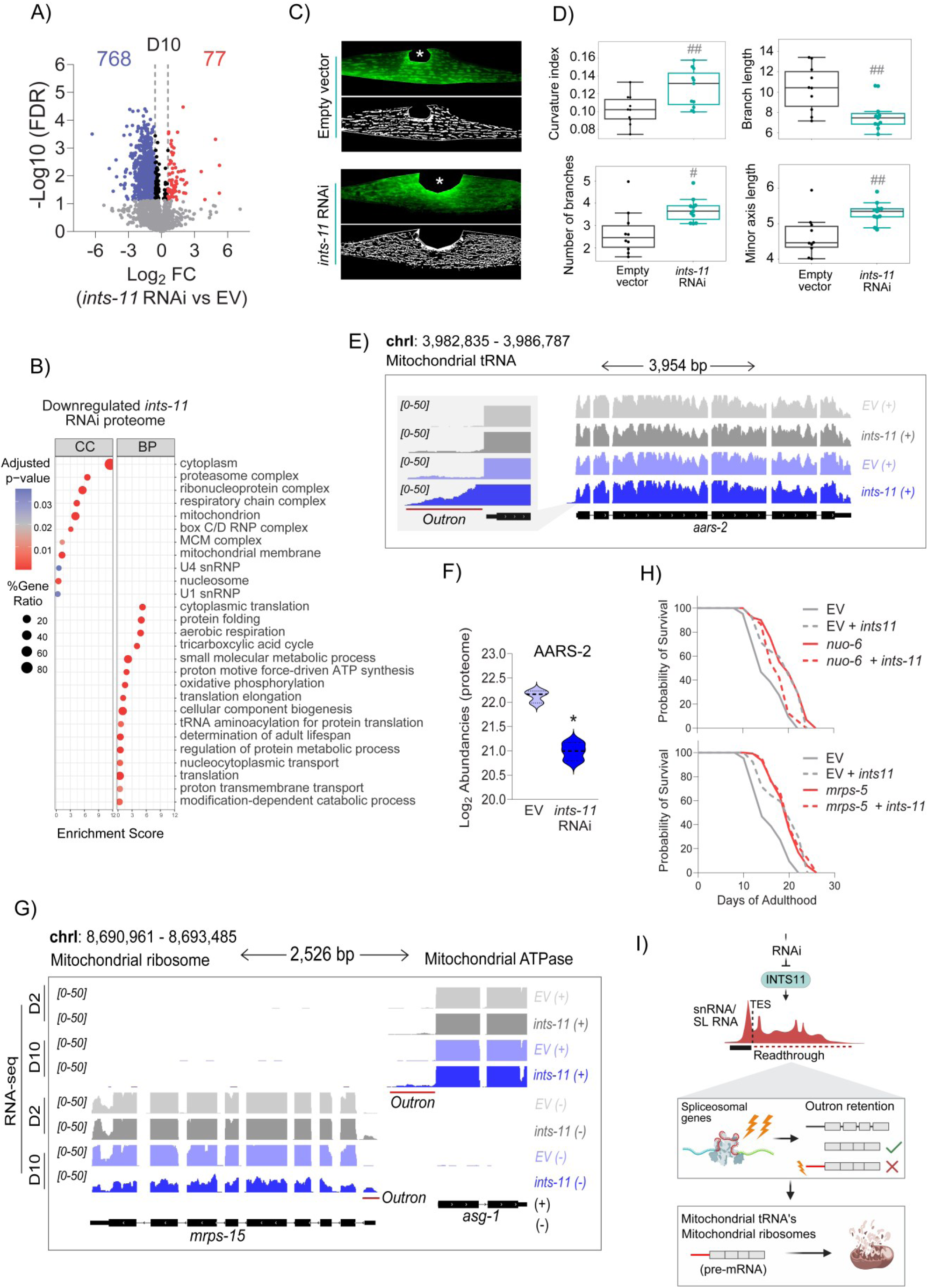
Proteomes of aged *ints-11-*depleted animals reflect defective spliceosomal function, negatively affecting mitochondrial protein expression linked to longevity. **A)** Volcano plot representing differentially expressed proteins in day-10 *ints-11* depleted animals. **B)** Gene ontology enrichment analysis (cellular compartment, left; biological process, right) of downregulated proteins in *ints-11* depleted animals. **C)** Fluorescence microscopy analysis of body-wall muscle mitochondria after 5-days of *ints-11* RNAi. **D)** Quantification of mitochondrial networks, implicating a ‘hyper-fused’ remodelling of mitochondria in *ints-11* silenced animals. **E)** Genome coverage tracks of a mitochondrial tRNA (*aars-*2, whereby depletion is known to extend lifespan) showing significant outron retention in *ints-11* depleted animals with no change in gene-body coverage. **F)** Protein abundance levels of AARS-2 are significantly reduced from *ints-11* RNAi in aged animals, suggesting outron retention impairs translation. **G)** Genome coverage tracks showing two nuclear-encoded mitochondrial genes (the ribosomal component *mrps-15,* and ATPase *asg-1*) required for mitochondrial function exhibit retained outrons in *ints-11* depleted animals. **H)** RNAi against two known longevity-promoting mitochondrial genes involved in electron transport chain function (*nuo-6*) and mitochondrial ribosome biogenesis (*mrps-5*). Combined silencing with *ints-11* RNAi is not additive to lifespan extension, implicating mitochondrial dysfunction in *ints-11*-mediated longevity axes. **I)** Model proposing how defective termination of sn/SL RNA genes impacts downstream splicing genes, leading to impaired splicing (outron removal) of mitochondrial tRNAs and ribosomes required for mitochondrial function.

Surprisingly, abundance of multiple mitochondrial (oxidative phosphorylation and membrane-related) proteins were also decreased in *ints-11* knockdown (Fig. S5C). To explore whether mitochondrial function was impacted upon loss of catalytic subunit of the Integrator, we validated mitochondrial integrity *in vivo* using a well-established transgenic reporter strain expressing GFP within body-wall muscle mitochondria (Fig. 5C). We found 5 days of *ints-11* silencing induced structural changes consistent with stress-adaptive remodelling, including increased mitochondrial curvature, branching and length (Fig. 5D). Together with the pronounced downregulation of mitochondrial membrane and respiratory proteins, these findings suggested activation of mitochondrial stress pathways such as the mitochondrial unfolded protein response (UPRmt) could potentially underpin longevity effects as has been well documented [80]. To test this, we assessed protein levels of the upstream UPRmt regulator ATFS-1, and monitored UPRmt activation using a p*hsp-6*::GFP reporter. However, neither ATFS-1 abundance nor *hsp-6*::GFP signal was increased following 5-days of *ints-11* RNAi (Fig. S5D–E), excluding canonical UPRmt activation as a contributor to longevity. Notably, recent studies indicate that active reproductive signalling is required for UPRmt engagement in *C. elegans* [81], consistent with the reduced progeny viability observed upon *ints-11* depletion (Fig. S1B).

We next considered whether mitochondrial dysfunction underlying longevity reflects impaired trans-splicing of mitochondrial genes. Consistent with this, mitochondrial tRNA-related proteins were among the most prominently downregulated in our proteomic dataset (Fig. 5B). A notable example was AARS-2 (human AARS2), which encodes the mitochondrial alanyl-tRNA synthetase required for mitochondrial protein synthesis [82]. Although *aars-2* mRNA levels were unchanged (Fig. 5E), AARS-2 protein abundance was markedly reduced following *ints-11* RNAi (Fig. 5F). Importantly, *aars-2* exhibited elevated outron-retaining reads (Fig. 5E), suggesting defective trans-splicing compromises productive protein output. Loss of *aars-2*, but not other mitochondrial or cytoplasmic tRNA genes, has previously been shown to extend lifespan in *C. elegans* [83]. Additional mitochondrial genes, including those required for mitochondrial ribosome assembly and electron transport chain function also displayed outron retention (Fig. 5G & S5F).

Mitochondrial dysfunction has previously been linked to lifespan extension in *C. elegans*. We therefore wondered whether the effect of *ints-11* mediated splicing dysfunction on mitochondrial genes was involved in longevity phenotypes. To test this we silenced *nuo-6* (NDUFB4) and *mrps-5* (MRPS5), core components of mitochondria’s complex I within the respiratory chain and mitochondrial ribosome biogenesis machinery, respectively. Both have previously been established to extend lifespan [84, 85]. Silencing either gene extended lifespan; however, no further lifespan extension was observed with *ints-11* RNAi (Fig. 5H), suggesting that mitochondrial dysfunction contributes to *ints-11* RNAi-induced longevity (Fig. 5I).

### microRNA responses are dispensable for *ints-11*-mediated lifespan extension

Previous work implicated Integrator in small RNA biogenesis in *C. elegans* [32, 48]. We therefore investigated whether perturbed small RNA levels might be involved in longevity phenotypes using small RNA-sequencing in day-10 depleted animals. We first investigated microRNAs, a major class of small RNAs which have previously been implicated in *C. elegans* lifespan regulation [86, 87] and were recently reported to depend on Integrator for RISC loading [30]. We observed that most mature microRNAs were upregulated upon depletion of *ints-11* (Fig. 6A). This was likely due to increased microRNA transcription because primary precursor transcripts were also elevated (Fig. 6B). Loss of microRNAs has been shown to reduce lifespan, so we wondered whether observed elevated microRNA levels might contribute to increased lifespan under *ints-11* knockdown. We utilised a mutant strain with auxin-inducible degron tagged to the endogenous loci of PASH-1 required for microRNA production [88]. PASH-1 depletion starting from young-adulthood led to rapid deteriorations in lifespan, as observed previously [89, 90] (PASH-1::AID auxin treated = 11 days median lifespan versus 18 days in wild-type auxin treated; Fig. 6C). However, we found that *ints-11* RNAi still extended lifespan in the absence of microRNA production, suggesting dispensability for microRNAs in *ints-11*-mediated longevity (Fig. 6C).

**Figure 6.**
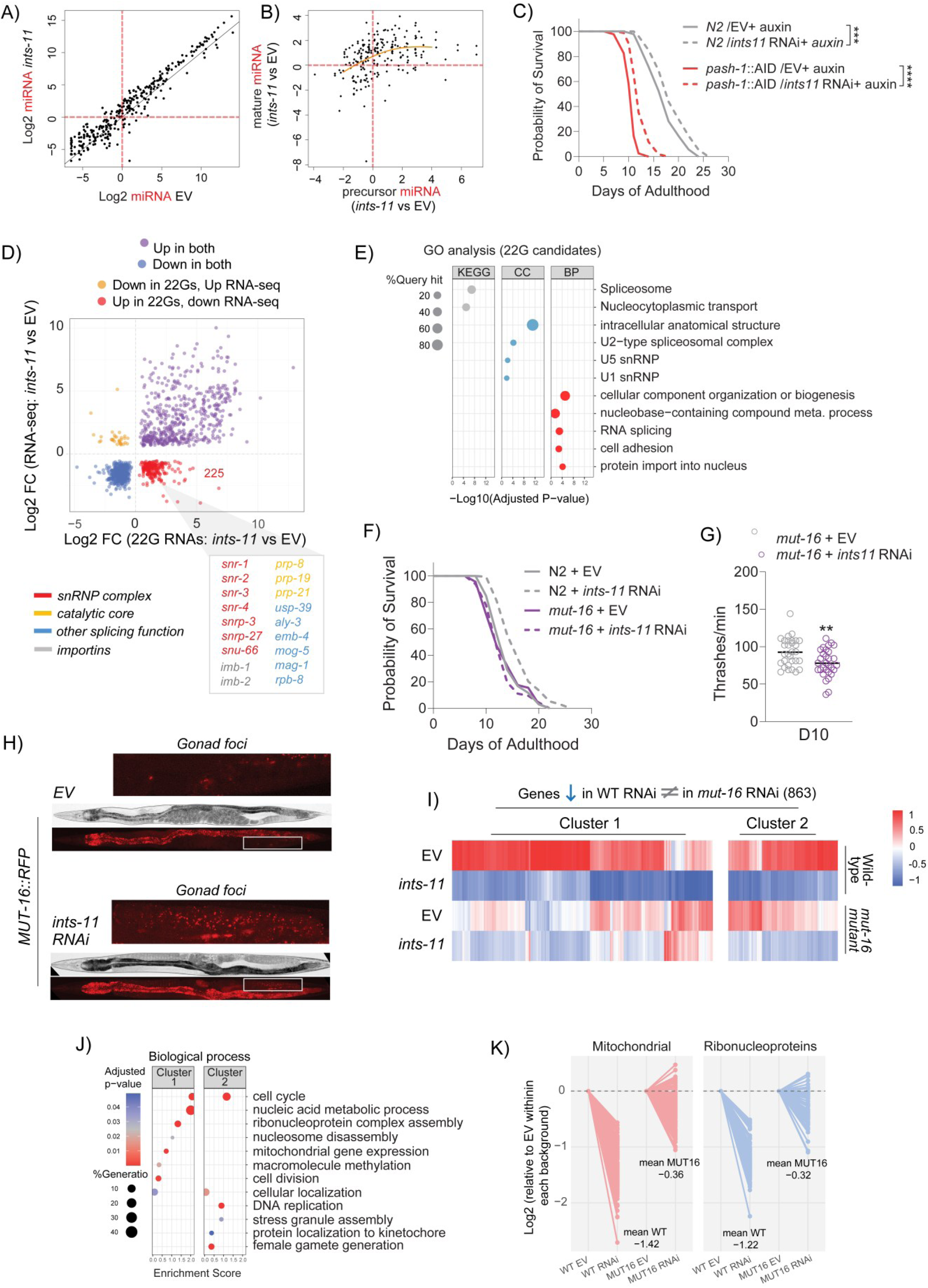
Endogenous siRNA biogenesis contributes to *ints-11*-mediated longevity and targets aberrantly processed spliceosomal genes in the gonad. **A)** Log2 scatter plot of mature micro-RNA levels in day-10 animals. **B)** Scatter plot of precursor *vs* mature micro-RNA levels in *ints-11* RNAi compared to empty vector controls. **C)** micro-RNA production is dispensable for the longevity effects of *ints-11* depletion. **D)** Quadrant plot of 22G RNA levels from small RNA-sequencing (x-axis) against the full-length target RNA. 225 represent as candidate endogenous siRNAs (increased 22G + decreased full RNA level, indicative of silencing). **E)** Gene ontology analysis identified candidate endogenous siRNAs as enriched for spliceosomal and nucleocytoplasmic transport genes. **F)** RNAi against *ints-11* within a *mut-16* deficient background (*pk710* allele) was unable to extend lifespan, implicating 22G RNA biogenesis as required for longevity effects. **G)** Healthspan attenuation is abolished from *ints-11* RNAi in *mut-16 (pk710)* deficient animals. **H)** Fluorescence microscopy of endogenously tagged MUT-16 (22G RNA biogenesis regulator). Silencing of *ints-11* for 2-days increased the number of MUT-16 foci within the gonad. **I)** Heatmap showing genes significantly downregulated in wild-type animals fed *ints-11* RNAi, where effects are reversed (cluster 1) or reduced (cluster 2) when performed in *mut-16(pk710)* mutants. **J)** Gene ontology enrichment analysis of gene sets reversed/reduced in *mut-16(pk710)* mutants under *ints-11* RNAi compared to wild-type RNAi. **K)** Representative line plot of mitochondrial and ribonucleoprotein genes downregulated from *ints-11* RNAi in wild-type animals, reversed in *mut-16(pk710)* mutants: abolishment of 22G RNA production restores expression levels of genes implicated in longevity axes from adulthood depletion of *ints-11*.

**Figure 7.**
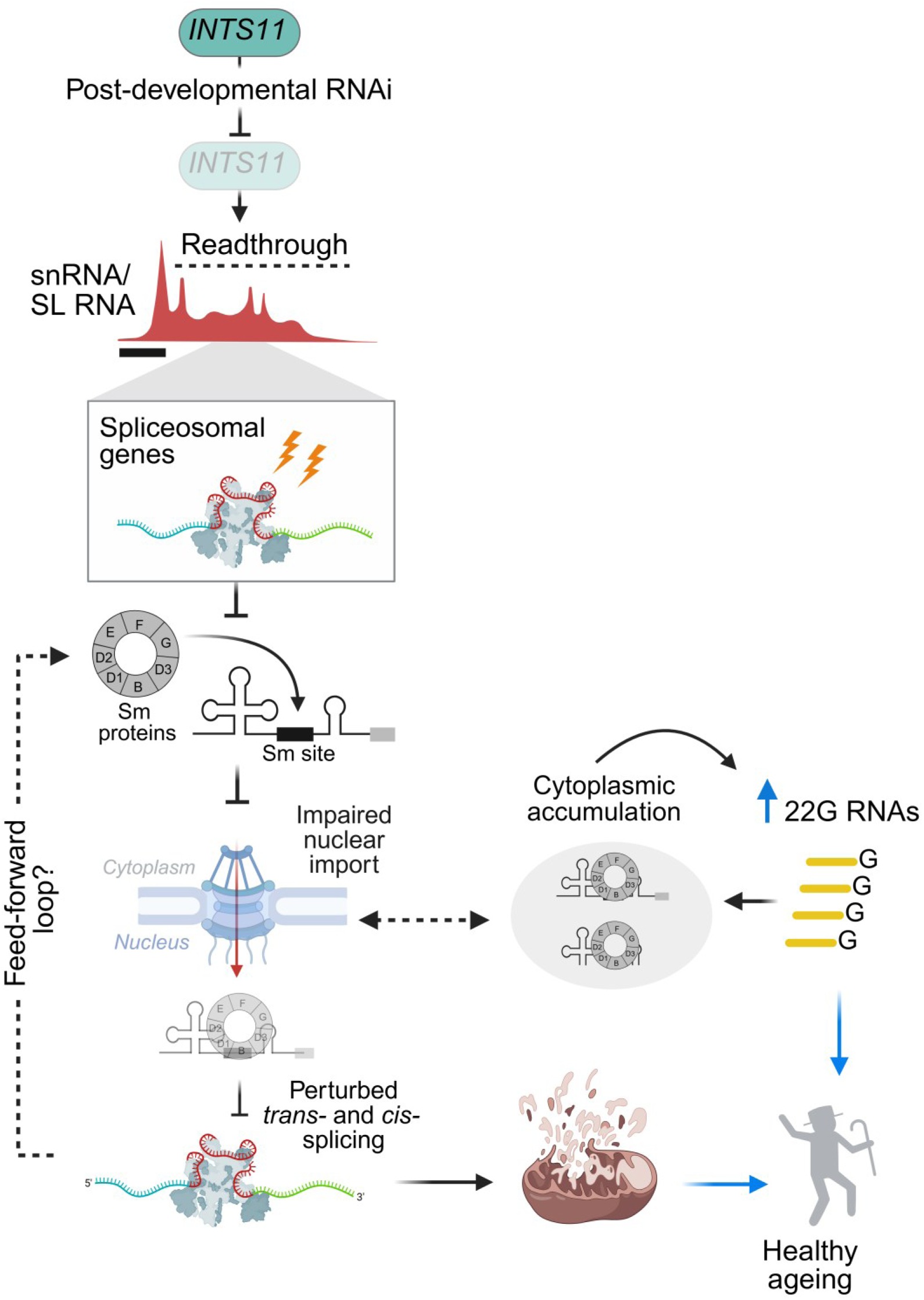
Proposed multi-modal mechanisms regulating lifespan extension from adulthood depletion of Integrator. Stark transcription termination defects of snRNA and SL RNA genes are evident within days of INTS11-targeted RNAi. Beyond impaired sn/SL RNA transcription, spliceosomal genes are preferentially impacted as they fall within extended readthrough intervals. This initiates a feed-forward cascade in which defective trans-splicing induces outron retention, disproportionately impairing spliceosomal components and the nuclear import machinery required to recycle snRNP and splice-leader complexes for subsequent splicing. We propose that accumulation of improperly processed transcripts in the cytoplasm engages 22G RNA pathways, which selectively silence genes enriched for spliceosomal and nuclear transport functions. As trans-splicing defects intensify with prolonged INTS11 depletion during ageing, outron retention extends to additional pathways, including mitochondrial genes. Ultimately, impaired trans-splicing, activation of 22G RNA biogenesis and mitochondrial dysfunction converge to drive lifespan extension following post-developmental INTS11 silencing.

### 22G RNA biogenesis is required for lifespan extension upon *ints-11* knockdown

Next, we examined whether small interfering RNAs played a role in Integrator-dependent lifespan extension. In *C. elegans*, RNA-dependent RNA polymerases generate 22-nucleotide, guanine-biased small RNAs (22G RNAs) that guide Argonaute-dependent silencing of complementary transcripts [90]. Notably, 22G RNAs have been implicated in lifespan extension induced by mitochondrial dysfunction [91]. To determine whether 22G pathways contribute to *ints-11*-mediated longevity, we mapped small RNA-seq reads to identify candidate 22G targets. Genes exhibiting increased 22G abundance coupled with reduced mRNA levels, consistent with 22G-dependent silencing, were classified as potential mRNA targets of 22G RNAs (n=255; Fig. 6D and supplemental table 5). KEGG pathway gene ontology analysis revealed strong enrichment for spliceosomal and nucleocytoplasmic transport pathways (Fig. 6E). These included multiple Sm/snRNP components (*snr-2, snr-3, snr-4, snrp-3, snrp-27, snu-66*), importins (*imb-1, imb-2*), and core splicing regulators (*prp-8, prp-19, prp-21, usp-39, aly-3*; Fig. 6D), many of which exhibit outron retention and reduced RNA and/or protein abundance.

We next tested whether 22G biogenesis is required for *ints-11*-induced longevity. MUT-16, a perinuclear scaffold essential for 22G amplification [92], is disrupted in *mut-16*(*pk710*) mutants, resulting in broad depletion of 22G RNAs. Strikingly, *ints-11* RNAi failed to extend lifespan in *mut-16*(*pk710*) animals (Fig. 6F). Moreover, the improved healthspan observed under *ints-11* RNAi was not only abolished but reversed in *mut-16*(*pk710*) mutants, indicating that 22G-mediated regulation is necessary to buffer physiological decline upon *ints-11* depletion (Fig. 6G). Importantly, although not shown here, we still observed the egg hatching defects in this background confirming that *ints-11* RNAi was still functional. Finally, imaging of an endogenously tagged MUT-16::RFP strain revealed increased MUT-16 foci within gonad arms after only 2-days of *ints-11* RNAi (Fig. 6H), consistent with activation of germline 22G pathways and supporting our earlier observed gonad-centric mechanism of lifespan regulation.

We next tested whether 22G RNAs contribute to longevity by silencing genes earlier implicated in lifespan extension. If so, loss of 22G RNAs should reverse *ints-11* RNAi-induced expression changes at these loci. To examine this, we performed RNA-seq in *mut-16*(*pk710*) mutants and applied a DESeq2 model incorporating genotype, treatment, and their interaction, enabling separation of baseline genotype effects from genotype-specific responses to *ints-11* RNAi. Genes were classified as reversed, attenuated, or unchanged in *mut-16*(*pk710*) based on effect size, direction, and interaction significance (FDR < 0.05, log2 fold-change ≥ 0.5). We identified 863 genes that were downregulated in wild type under *ints-11* RNAi but exhibited increased expression, attenuation, or no change in *mut-16*(*pk710*) animals following *ints-11* RNAi (Fig. 6I). Gene ontology analysis of this reversed set revealed significant enrichment for ribonucleoprotein and mitochondrial pathways (Fig. 6J). Notably, mitochondrial genes showed a mean log2 expression reversal of 1.06, while ribonucleoprotein assembly genes exhibited a mean reversal of 0.90 (Fig. 6K).

## Discussion

Age-related functional decline is driven, in part, by pleiotropic alterations in Pol II transcription [6]. Despite this, contribution of the Integrator complex, a central regulator of Pol II transcription, remains largely unexplored in shaping the pace of ageing. Here, we identify the Integrator catalytic subunit INTS11 as a post-developmental determinant of longevity. Depletion of INTS11 disrupts snRNA transcriptional termination and trans-splicing, impairing outron removal in spliceosomal, nuclear import, and mitochondrial genes, acting to limit protein expression. Defective RNA processing engages endogenous siRNA pathways and induces a decline in mitochondrial function, collectively reprogramming cellular homeostasis toward lifespan extension.

### Conserved and divergent genomic functions of Integrator

In mammals, Integrator is enriched at promoter-proximal paused Pol II at coding and non-coding genes. Depletion of Integrator subunits leads to termination defects at snRNAs and upregulation of enhancer [9] and bidirectional-promoter antisense transcription in mammalian cells [20], consistent with the idea that it suppresses unwanted cryptic transcription. We mapped INTS11 genomic occupancy for the first time in *C. elegans* and revealed INTS-11 bound promoters/TSSs, but that INTS-11 was also enriched at TESs and within intragenic regions, suggesting a somewhat distinct binding profile to mammals. Nevertheless, we observed conserved activity at non-coding regulatory loci: INTS11 extensively bound enhancers, and *ints-11* RNAi caused enhancer de-repression and widespread termination defects at snRNA consistent with a requirement for Integrator’s endonucleolytic activity. We find that INTS-11 occupancy does not convincingly correlate with differential gene expression in RNA-seq data, as many unbound genes also show significant expression changes, likely due to transcription-termination defects at noncoding loci. Comparative ChIP-seq and RNA-seq analyses are however limited by the fact binding profiles represent only a snapshot of INTS-11 occupancy, that binding may differ in aged animals, and that immunoprecipitation efficiency and recovery is variable across genes.

Despite these limitations, we found that INTS-11 depletion leads to a reduction in sense-strand transcription downstream of promoters targeted by Integrator, consistent with impaired productive elongation similar to what has been reported in higher mammalian systems. For example, ∼50% of differentially expressed transcripts were previously shown to be downregulated from depletion of INTS9 and INTS11 in human cells [27]. In contrast, studies in *Drosophila* have reported a striking upregulation of protein-coding genes (>80% of differentially expressed genes) upon INTS9 depletion [8, 18]. It was proposed that Integrator supports productive elongation by mediating release of elongation-incompetent Pol II complexes and mediates recycling and initiation of functional Pol II. Our data support that the function of Integrator in elongation is conserved in *C. elegans* despite the lack of wide spread Pol II pausing at promoters [93] and the absence of a conserved NELF complex [62]. In fact, the lack of NELF might help explain the somewhat contradictory finding for limited promoter-proximal attenuation and suppression of PROMPT-like transcripts: in vertebrates, NELF stabilises promoter-proximal paused Pol II and functions together with Integrator [61, 94]. However, gene specific Integrator-dependent effects have been recently proposed to be mediated via Integrator’s interaction distinct transcription factors [36, 60]. Our data therefore raise the possibility that, although NELF-dependent pausing architectures are absent in nematodes, Integrator-mediated quality control of Pol II complexes to ensure productive elongation is evolutionarily conserved.

Lastly, an important feature of our work to highlight, is that *C. elegans* possess a sub-type of non-coding snRNA, not present in higher mammals, that exhibits transcription-termination defects. These are the SL RNA (both SL1 and SL2 RNAs) that are required for trans-splicing function [70, 73, 74, 78]. *C. elegans*-specific requirements for Integrator have been reported previously, including roles in piRNA biogenesis [32, 48] and for transcription-termination of SL RNAs throughout development [19]. Here, we show that Integrator endonucleolytic activity is required for trans-splicing function throughout the genome, adding to the current *C. elegans*-specific repertoire of Integrator functions.

### Defective trans-splicing regulates lifespan extension

We provide the first evidence that direct post-developmental inhibition of trans-splicing can extend *C. elegans* lifespan, and that perturbed trans-splicing is in fact the early, causative mechanism by which loss of INTS11 extends lifespan. We propose that early trans-splicing loss initiates a positive-feedback loop: outron retention preferentially affects spliceosomal and nuclear import genes required for mature SL RNP biogenesis and nuclear re-import. The resulting decline in protein levels, consistent with the role of trans-splicing in enhancing translational efficiency [79], limits nuclear SL RNP availability, further constraining trans-splicing of these same genes. As trans-splicing capacity progressively deteriorates, defects extend into wider biological networks such as nuclear-encoded mitochondrial genes that then drive lifespan extension, a well-established biological phenomenon in *C. elegans* [95] and higher mammals [96].

We also found that endogenous siRNA production was required for INTS11-mediated longevity. siRNAs mapped predominately to spliceosomal and nucleocytoplasmic genes, suggesting a role in mitigating mis-processed spliceosomal RNA. Intriguingly, previous data has shown INTS4 and INTS11 depletion in HeLa cells induces aberrant snRNA 3’ end extension that impairs nuclear export and disrupts Cajal body integrity in human cells [97]. Critically, a key consequence was the accumulation of cytoplasmic Sm/snRNP components largely devoid of mature snRNAs, reported to limit snRNP maturation and subsequent nuclear re-import. In addition to reflecting conserved defects in snRNP biogenesis, the cytoplasmic accumulation of these factors supports the notion that endogenous siRNAs in *C. elegans* may uphold lifespan/ healthspan effects by promoting degradation of proteotoxic spliceosomal components. Consistent with this model, the healthspan benefits of INTS11 RNAi, such as enhanced neuromuscular activity, are not only lost but become detrimental in 22G RNA-deficient backgrounds.

### Trans-splicing: an unexplored regulator of the proteome in ageing

A central finding of our study is that impaired trans-splicing of pre-mRNAs is associated with downregulation of the corresponding proteins. This aligns with previous biochemical evidence showing that trans-splicing shortens native 5′ sequences and removes upstream AUGs, thereby enhancing translation initiation efficiency in *C. elegans* and increasing protein production [79].

A recurring theme in multi-omics studies, particularly in *C. elegans*, is the poor correlation between mRNA and protein abundance [98, 99]. While technical factors (e.g. poly(A)-based mRNA enrichment, which limits insight beyond transcript-level up/downregulation)) may contribute to this discrepancy, our data suggest an additional explanation: alterations in trans-splicing may represent an overlooked layer of transcriptional regulation in ageing. We observe multiple cases in which RNA-seq coverage remains stable, yet outron retention is associated with marked reductions in protein abundance (e.g. AARS-2). Furthermore, many genetic interventions that extend lifespan or healthspan in *C. elegans* fail to translate effectively to higher eukaryotes. One possible explanation is that trans-splicing-mediated regulation contributes to these phenotypes in nematodes but is absent in mammals. Indeed, it is plausible that some well-established lifespan-extending mutations act, at least in part, through modulation of trans-splicing efficiency. Elucidating this mechanism would clarify this translational gap and help future studies in *C. elegans* ageing studies.

### Integrator as a pleiotropic transcriptional complex

The AP theory of ageing posits that genes promoting early-life fitness can have detrimental effects later in life [54]. As the force of natural selection declines after reproduction, alleles that enhance early survival are retained despite late-life costs, limiting evolutionary pressure to optimise gene function for healthy ageing [52]. In this context, Integrator might be viewed as exhibiting features consistent with AP, functioning as a transcriptional regulator that is fundamental in early-life but deleterious in older age. Indeed, our finding that silencing INTS11 from day-4 of adulthood prolongs lifespan is a classical genetic illustration of AP.

However, an important and often underappreciated consideration is that longevity arising from depletion of essential genes may reflect secondary adaptive responses rather than the direct effects of wild-type gene function. In our case, silencing INTS11 in adulthood extends lifespan, yet the pro-longevity effects stem from downstream consequences of defective snRNA transcription/ trans-splicing rather than from any intrinsically deleterious property of INTS11 itself. Thus, wild-type INTS11 is unlikely to actively promote ageing, instead, loss of its essential transcriptional role initiates a compensatory cascade that enhances late-life healthspan. These distinctions warrant caution when claiming AP, as the ageing phenotype may arise indirectly from stress-induced adaptive reprogramming rather than from direct late-life costs of wild-type gene function.

Further arguing against a straightforward AP model, Integrator’s impact on lifespan was markedly subunit- and tissue-specific. Subunits associated with catalytic or recruitment functions, including INTS4/11/13 and the PP2A-positioning components INTS2/5/6/8, primarily regulate RNA processing and transcriptional termination rather than maintaining core complex architecture. Partial attenuation of these activities may therefore permit adaptive transcriptional reprogramming to dominate, promoting longevity. In contrast, INTS1 and INTS7 reside within the structural core of Integrator and form key interfaces required for complex stability and the INTS3–INTS7 interaction that facilitates release from Pol II [25]. Loss of these subunits may more broadly destabilise Integrator architecture, pushing RNA-processing dysfunction beyond a hormetic threshold and accelerating ageing. The pronounced negative phenotype observed upon INTS3 depletion is further consistent with its additional roles in DNA damage repair and genome stability pathways [100], potentially reducing tolerance to Integrator disruption. Future biochemical work will be essential to address this gap. For example, complex purification coupled to quantitative mass spectrometry could determine the extent to which depletion of distinct modules disrupts full Integrator assembly and stability. In parallel, defining how depletion of multiple subunits spanning different modules shapes adult healthspan will help pinpoint the mechanisms underlying the divergent phenotypic effects observed here. Taken together, our data indicate that lifespan extension arises not from a deleterious late-life action of Integrator itself, but from the systemic adaptive responses triggered by partial disruption of its RNA-processing functions.

### Integrator as a putative target for human ageing

An important translational distinction is that trans-splicing does not occur in higher mammals, nor do they possess endogenous RNA-dependent RNA polymerase pathways that generate 22G RNAs. These differences complicate direct extrapolation of our findings to humans and raise the question of whether selective attenuation of Integrator subunits could beneficially modulate ageing in mammalian systems.

Defective snRNA transcription termination in human cells impairs snRNP biogenesis and nuclear re-import [97], providing a mechanistic parallel to the RNA-processing defects observed in *C. elegans*. Nonetheless, our data indicate that worm longevity is driven primarily by disruption of SL RNP-dependent trans-splicing rather than snRNP-mediated alternative splicing. In organisms lacking trans-splicing, impaired snRNA processing may instead influence ageing through altered spliceosome function and isoform selection. Such changes could reshape downstream pathways, such as mitochondrial bioenergetics, to impact lifespan in mammals. However, in the absence of RNA-surveillance mechanisms analogous to 22G RNA pathways, cytoplasmic accumulation of misprocessed snRNP components may constrain the capacity for ageing attenuation, an effect we reveal here in *C. elegans*. One possibility is that alternative RNA-regulatory pathways, such as microRNA-mediated control, fulfil a compensatory role in mammals. Notably, Integrator has recently been implicated in regulating microRNA abundance and RISC loading in human cells [30], suggesting a potential parallel mechanism linking RNA processing to post-transcriptional homeostasis.

Critically, the effects of late-life manipulation of Integrator subunits have not been rigorously examined in primary human cells capable of replicative senescence. Addressing this gap will be essential to determine whether Integrator represents a viable target for attenuating age-related decline in humans.

## Data availability

RNA-seq, small RNA-seq and ChIP-seq data have been deposited in NCBI’s Gene Expression Omnibus and are accessible through GEO Series accession numbers GSE325594, GSE325598 and GSE325597. Proteomics data have been deposited to ProteomeXchange with the identifier PXD076576.

## Acknowledgements

We thank members of the Vasilieva lab and Peter Sarkies for helpful discussions, advice, and valuable comments on the manuscript. This work was supported by the Wellcome Trust Senior Fellowship and BBSRC grants to L.V. (WT106994/Z/15/Z, BB/Y00194X/1 and BB/Y004590/1), the Wellcome Trust Investigator Award to P.S. (227739/Z/23/Z). JC was supported by Project PID2024-161136NB-I00, financed by the Spanish Ministry of Science, Innovation and Universities (MICIU).

## Author contributions statement

L.S., P.S. and L.V. conceived and designed the experiments. L.S. performed all experiments with the exception of small RNA-seq analyses that was performed by P.S. P.S. provided advice, contributed to the design of some of the experiments related to small RNAs and *C. elegans* genetics and performed analyses of small RNA-seq data and mRNA targets predictions. M.F. contributed to proteomics analyses and discussions. J.C. generated INTS-11-FLAG and provided several RNAi’s used in this study. Y.J. provided RNAi for the factors screened in this study. K.K. contributed to setting up ChIP-seq protocol in *C. elegans* and bioinformatics analyses. L.S., P.S. and L.V. prepared the figures and wrote the manuscript. All authors read and commented on the manuscript.

## Competing interests statement

The authors declare no competing interests.

## Materials and methods

*C. elegans* maintenance. Unless otherwise stated, strains were maintained at 20°C on standard nematode growth media (NGM) plates with OP50 as a food source as previously described [101]. All strains were purchased from the *C. elegans* genetic centre which is funded by the NIH Office of Research Infrastructure Programs (P40 OD010440). The strains used in this study were as follows: N2 (Wild-type Bristol isolate), CB4037 *glp-1(e2141)*, CB5600 *(ccIs4251)[Pmyo-3::mitoGFP]* I; *him-8(e1489)* IV., MLC1065 *pash-1(luc71)[pash-1::2xGGSG::3xFLAG::AID*::myc]* I; *ieSi57* II*; unc-119(ed3)* III*; ieSi38* IV., NL1810 *mut-16 (pk710)* I., SJ4100 *(zcls13)[hsp-6p::GFP + lin-15(+)]* V., AMJ345 *(jamSi2) II; rde-1(ne219)* V., CYA13 *(frSi17) II; rde-1(ne300)* V; *ldrIs1*., DCL569 *(mkcSi13) II; rde-1(mkc36)* V., IG1846 *(frSi21)* II; *frIs7 IV; rde-1(ne300)* V., RA382 *(rdIs7) I; unc-119(ed3) III; rde-1(ne219)* V., WM118 *rde-1(ne300)* V; *neIs9* X., XE1375 *(wpIs36)* I; *wpSi1 II; eri-1(mg366)* IV; *rde-1(ne219)* V; *lin-15B(n744)* X., XE1474 *(wpSi6)* II; *eri-1(mg366)* IV; *rde-1(ne219)* V; *lin-15B(n744)* X., XE1581 *(wpSi10)* II; *eri-1(mg366)* IV; *rde-1(ne219)* V; *lin-15B(n744)* X., XE1582 *(wpSi11)* II; *eri-1(mg366)* IV; *rde-1(ne219)* V; *lin-15B(n744)* X., SHG2030 *mut-16 (ust366 [mCherry::mut-16])* I., LVA01 (*syb10825 [ints-11p::GFP]*).

### RNA interference

Adult worms were fed HT115 E. coli carrying either an empty vector (L4440) or vectors expressing double-stranded RNAi. All constructs were obtained from the Ahringer library and sequence verified by Sanger sequencing before use. RNAi bacteria were grown on LB plates supplemented with 100µg/mL Carbenicillin + 12.5 µg/mL Tetracycline overnight at 37°C. Single colonies were picked and grown in LB broth supplemented with 100µg/mL Ampicillin for approximately 6-7 hours at 37°C with shaking at 200RPM. OD-normalised bacteria were seeded onto dry RNAi NGM plates (50 mM NaCl, 0.25% (w/v) bacteriological peptone, 1.7% (w/v) agar, 1 mM CaCl2, 1 mM MgSO4, 25 mM KH2PO4 (pH 6), 12.9 μM cholesterol, supplemented with 100µg/mL Carbenicillin and 1mM IPTG)) and allowed to grow for 2-days at room temperature to allow dsRNA induction. For reproducibility, RNAi plates were made in batch and stored at 4°C until use for up to 3-weeks. All experiments contained a positive control of pop-1 RNAi (induces strong dead egg phenotype) to ensure robust RNAi efficiency.

### Tissue-specific RNAi

Strains deficient in *rde-1* (spanning *ne219* and *ne300* alleles) were used, whereby rde-1 has been restored back under a tissue-specific promoter [102, 103]. Ahringer RNAi constructs were then used for feeding in these backgrounds, where RNAi was isolated to the respective tissue under rde-1 promoter-driven restoration. These strains have been generated previously [102, 103] and were obtained from the *C. elegans* genetic centre (see maintenance section above). All RNAi feeding was started from the L4 stage.

### Combined RNAi feeding

For combining dsRNA feeding clones, bacteria were grown as described above. After 6-7 hours of growth at 37°C, bacterial concentrations were assessed by OD. To first generate single RNAi cultures, all clones were normalised to equal OD’s using empty vector (L4440) as a dilutant. For combined RNAi, clones were then mixed 1:1 to generate mixed RNAi pools of two dsRNA clones. Bacteria were seeded onto RNAi plates and allowed to grow for 2-days at room temperature.

### Auxin-inducible degradation

A 400 mM auxin stock solution was prepared by dissolving indole-3-acetic acid in ethanol, followed by filter sterilisation and storage at 4°C in light-protected tubes as previously described [104]. NGM agar (supplemented with 100µg/mL Carbenicillin and 1mM IPTG) was prepared and divided into two equal portions. Auxin stock was added to one portion to achieve a final concentration of 0.15 mM, while the second portion received an equivalent volume of filter-sterilised 100% ethanol to serve as a solvent control. Due to light sensitivity, both auxin-containing and control plates were protected from light during overnight drying using foil or a black covering and subsequently stored at 4°C. Plates were removed from 4°C and left covered at room temperature the night before seeding with HT115 bacterial cultures. For lifespans, animals were grown on regular OP50-seeded NGM plates from egg hatching to L4 larval stages, and then transferred onto either ethanol- or auxin-treated RNAi plates as young adults.

### Lifespan assays

*C. elegans* were age-synchronised by egg laying and maintained on standard 90mm NGM plates seeded with 1mL of OP50 bacteria from hatching until L4 (young adult) larval stage. For RNAi experiments, animals were then transferred to respective RNAi plates (55mm) and transferred/ counted on fresh plates every 2-days. Animals were classified as dead if they did not display any response to gentle prodding of the head three times. Animals with vulval protrusion or that were missing/ dried out on the edge of plates were censored.

### Motility healthspan assays

To assess animal healthspan, we used manual thrashing performance as a robust proxy for neuromuscular health known to correlate strongly with mortality rate. On the day of assay, individual worms were picked into 10µL M9 buffer (3g KH2PO4, 6g Na2HPO4, 5g NaCl, 1 mL 1M MgSO4 per litre) on glass slides. Body-bend counts were done manually, with each full bend left and right marking a count. Counts were measured over 20-seconds and displayed as thrashes per-minute.

### Egg laying assays

.At the required age, ten individual adults per condition were placed onto 55mm plates and allowed to lay eggs for 6-hours. Adults were then removed and plates containing eggs were placed at 20°C until eggs developed to L4 stage. Plates were then moved to 4°C for 2-hours to aid manual counting of hatched L4 adults per plate.

### Heat stress assays

To test overall organismal robustness, animals were fed either control (L4440) or dsRNA bacteria starting from the L4 stage, maintained on respective plates with transferring every 2-days until day-10 of adulthood, and then shifted to 37°C for thermotolerance assays. Plates were placed at 37°C and then individuals subject to survival analysis every hour thereafter. Animals were scored as dead if they did not respond to gentle prodding of the head more than three separate times.

### Microscopy analysis

All imaging was performed on a Leica Thunder fitted with an sCMOS Hamamatsu Orca Fusion Digital C14440-20UP camera, with images captured using either 40X HC PL APO Dry NA 0.95 (OM42) or 63X HC PL APO CS2 water automated immersion objectives and a cooled pE-800 LED Light Engine. Worms were grown to the desired stage and then manually picked into 18µL of M9 buffer on glass slides. Worms were gently immobilised by cover slips and immediately placed on stage for imaging. Mitochondrial network analysis was performed with MitoSegNet [105] using default parameters optimised for the specific strain utilised in this study. Network quantification was subsequently performed using the MitoA package within the MitoSegNet pipeline in order to generate quantitative morphological analysis of masked mitochondrial networks. All other image analysis was performed in ImageJ.

### Western blotting

Worm pellets were collected and lysed in RIPA buffer supplemented with protease inhibitors. Lysates were boiled for 5 min and clarified by centrifugation. Equal amounts of total protein were resolved on 4-12% gradient SDS-PAGE gels and transferred to PVDF membranes by wet transfer (2h at 4°C). Membranes were blocked in 5% non-fat milk in TBST and incubated overnight at 4°C with anti-FLAG-HRP-conjugated antibody (Merck, A8592) at 1: 2,000 dilution. Following washes in TBST, signals were detected using chemiluminescence and imaged using a LI-COR Odyssey imaging system.

### RNA isolation, library preparation and sequencing

#### Sample collection

.Animals were grown on OP50 bacteria until L4 stages, and then shifted to either L4440 or dsRNA-seeded plates until the desired age. As we performed Ribodepleted RNA-sequencing, animals were manually picked from bacterial seeded plates to unseeded NGM plates for 20-minutes to remove as many bacteria from within the gut and on the exterior cuticle. Given Ribodepleted RNA-sequencing methods do not utilise mRNA enrichment for library preparation, we found this approach (as opposed to washing worms straight from plates into tubes) assisted with limiting the extraction and amplification of bacterial RNA that contaminates read allocation during sequencing. Worms (∼80 per condition) were then picked into 1.5mL low-bind Eppendorf’s containing 30µL of M9 buffer, washed six times with 1mL M9 buffer, and the final supernatant removed as close to the worm pellet as possible. Samples were mixed with 500µL of TRIzol reagent, snap frozen in liquid nitrogen and stored at -70°C until further processing.

#### RNA Extraction

RNA was isolated by performing three freeze-thaw cycles, with vortexing for 10-seconds in between. Samples were then mixed with 100µL of chloroform with pulse vortexing for 10-seconds before a 20-minute centrifugation at 15,000g and 4°C in phase-separating tubes to avoid phenol contamination. The aqueous solution was placed into a fresh tube, incubated with 100% isopropanol for 5-minutes with mixing and then centrifuged for 15-minutes at 15,000g and 4°C to pellet RNA. Pellets were then washed two-times with 75% ethanol, and evaporated to remove residual contaminants before re-suspension in DNAse free pure water. All purified RNA was assessed via Qubit High Sensitivity RNA for quantities and Bioanalyzer for RNA integrity before downstream processing.

#### RNA processing

Equal quantities of RNA were taken and subject to spike-in using RNA from Schizosaccharomyces pombe at a constant ratio of 5% across experiments. Spiked-in samples were then subject to DNAse treatment to remove contaminating DNA, immediately precipitated to quench reaction enzymes, washed in 75% ethanol two-times and evaporated for resuspension in ultra-pure water. Samples were then normalised to a total of 500ng RNA in 14µL using Qubit for Ribodepletion using the *C. elegans* RiboPOOL from siTOOLs as per the manufacturer’s instructions. Ribodepleted RNA was purified via overnight precipitation at -20°C, centrifuged for 30-minutes at 15,000g and 4°C the following day, washed and evaporated dry. All samples were check for Ribodepletion efficiency using Bioanalyzer compared to pre-depleted RNA. RNA quantities were checked again via Qubit, and samples were normalised for library preparation using the NEBNext Ultra II Directional RNA Library Prep Kit for Illumina as per the manufacturer’s instructions and sequenced as paired-end 40bp reads on an Illumina NextSeq 500.

### Bioinformatic analysis of RNA-seq data

RNA-seq libraries (paired-end, 40 bp, reverse-stranded) were processed using a custom Python-based pipeline integrating fastp, STAR, samtools, bedtools and deepTools. Libraries contained *Schizosaccharomyces pombe* spike-in RNA; however, spike-in normalisation was not applied due to variability in gut bacterial content (and thus amplification bias) across samples. This is an important note to consider for future Ribo-depleted total RNA-seq in *C. elegans,* whereby variations in this third unknown variable limit capacity for accurate spike-in normalisation. Downstream analyses were therefore normalised by routine library size factors.

Raw FASTQ files were quality filtered and adapter trimmed using fastp (-l 30, --detect_adapter_for_pe, -w 4). Trimmed reads were aligned using STAR to a concatenated genome containing *C. elegans* (WBcel235) and *S. pombe* references (--runThreadN 18, --readFilesCommand zcat, --alignIntronMin 10, --alignIntronMax 5000, --alignMatesGapMax 10000, --outSAMtype BAM Unsorted, --outReadsUnmapped Fastx). BAM files were coordinate-sorted and indexed using samtools (samtools sort -@ 4; samtools index -@ 4). Reads were separated with samtools, then split by organism using samtools idxstats and samtools view. Paired reads with insert sizes exceeding 2 kb were removed using a custom filtering step. Uniquely mapped reads were extracted samtools view -q 255 and re-indexed. For visualisation, strand-specific coverage tracks were generated using deepTools bamCoverage (--filterRNAstrand forward/reverse, -bs 1, -p 6) with scaling based on library size. Coverage files were generated separately for each strand and exported as bigWig files.

Gene-level counts were generated using featureCounts against the WBcel235 GTF annotation. Differential expression analysis was performed in R using DESeq2 [106] with default median-of-ratios normalisation. Genes were considered significantly differentially expressed if adjusted P values were ≤ 0.05 and absolute log2 fold-changes were ≥ 1.

### ChIP-seq library preparation

Samples and buffers were prepared as described in [107], with optimisations. We utilised an endogenous *ints-11* FLAG-tagged strain (JCP643 *ints-11*(*jcp31*)[*ints-11*::3xFLAG])) for INTS11 ChIP-seq. Per replicate, approximately 400,000 age-synchronised worms (L1 stage after bleaching) were spread across thirty 150mm petri plates containing standard NGM seeded with 3mL of OP50 bacteria. Animals were grown to day-1 of adulthood, washed from plates with M9 buffer into 50mL falcon tubes, washed a further five times to remove bacterial debris and then frozen at -70°C. Pellets were then disrupted in a freezer mill (SPEX SamplePrep) in liquid nitrogen to generate 5mL of lysed worm powder per replicate. Lysed worm powder was then cross-linked in warm PBS supplemented with formaldehyde (1.1% final) and PMSF (1% final) in a total volume of 40mL. Tubes were placed on a rotator at room temperature for 10-minutes, and then quenched with glycine to a final concentration of 125mM. Cross-linked samples were then washed with 10mL of ice-cold PBS supplemented with protease inhibitors for a total of three washes with centrifugation at 4000g and 4°C. Pellets were resuspended in ice-cold B-ChIP-L0 buffer for chromatin shearing.

Approximately 1.7mL of sample was added to a 15-mL polymethylpentene Bioruptor falcon tube and mixed with 200µL of Bioruptor sonication beads. Samples were topped up to 2mL final with ice-cold B-ChIP-L0 buffer for optimal shearing conditions. Samples were manually vortexed in a cold room for 30-seconds, and rested on ice for 1-minute for a total of three times. Chromatin shearing was then performed using a Bioruptor Pico sonicator with maximum settings using 30-second on/ 90-seconds off for 8 cycles at a time. Samples were rested on ice for 1-minute in between, and briefly vortexed to re-homogenise samples. Sonication was repeated in this manner for a total of 72 cycles to achieve fragment sizes of approximately 150 – 300bp. Samples were centrifuged two times for 5-minutes at 20,000g and 4°C, and two 50µL aliquots of sheared solution were stored at -20°C to test for DNA fragment size (using Bioanalyzer) and later protein pull-down efficiency (via western blot). The remaining cleared supernatant was added to 1.5mL low-bind tubes and incubated with 10µL Anti-FLAG M2 Magnetic Beads (Merck, M8823) prepared in B-ChIP-BL buffer and rotated at 4°C for 4-hours. Samples were then placed on a magnetic rack for 1-minute and a further 50µL aliquot was stored at -20°C as the protein flow-through. Supernatants were removed and beads were carefully washed with 1.5mL ice-cold B-ChIP-Lys buffer and rotated for 5-minutes at 4°C – this step was repeated three times. Samples were then washed a further two times with B-ChIP-Lys buffer (now containing 200mM NaCl). A further three washes were then performed with ice-cold B-ChIP-W buffer, and three more times with 1.2mL of ice-cold TE (pH 8.0) containing 50mM NaCl. De-crosslinking was performed using 280µL of B-ChIP-EL buffer supplemented with 10µL of 10 mg/ml RNase A and incubated at 37°C for 1-hour with 800RPM shaking. 10µL of 10 mg/ml proteinase K was added and incubated for 2-hours at 55°C and 800RPM. Samples were then incubated overnight at 65°C with shaking, and purified using a ChIP DNA cleanup kit. Final fragment sizes were checked using Bioanalyzer, and library preparation done using the NEBNext Ultra II DNA Library Prep Kit for Illumina.

### Bioinformatic analysis of ChIP-seq data

ChIP-seq peak calling of INTS-11::3xFLAG immunoprecipitated samples was performed using MACS2 on aligned BAM files against background input samples. Peaks were identified using --qvalue 0.01, --nomodel, --extsize 250, and --keep-dup auto. This yielded 7,276 significant peaks used for downstream analyses.

The *C. elegans* annotation file (WBcel235) does not contain genomic maps of putative promoters and enhancers, for example, two features with well-known Integrator binding in mammals. We thus implemented a two-step approach to improve the accuracy of genomic feature binding. Firstly, we mapped peaks against a recently published annotation file by the Ahringer laboratory containing >42,000 regulatory elements [59]. Peak annotation was performed using a custom pipeline implemented in bash and Python using bedtools intersect -wo. For peaks overlapping multiple regulatory intervals, a single assignment was made based on maximal base-pair overlap from the MACS2 narrowPeak file. Peaks not assigned to Ahringer regulatory elements were subsequently intersected with the WormBase GTF annotation (WBcel235, release 113) using bedtools intersect -wo, and assigned to the gene/ feature with the greatest overlap. Gene-associated peaks were classified as TSS, TES, exonic or intronic based on overlap with corresponding GTF features; peaks lacking gene overlap were classified as intergenic. Intronic peaks were additionally evaluated using bedtools window -w 300 to identify proximity to opposite-strand TSS/TES regions, given the operonic nature of the *C. elegans* genome. Genes with at least one uniquely assigned peak were defined as INTS11-bound.

For metagene analyses, recruited genes with transcription start sites located within 300 bp of another same-strand gene were excluded to reduce tandem gene artefacts. For signal visualization, replicate ChIP-input normalized bigWig files were averaged using deepTools bigwigCompare --operation mean --binSize 1. Metagene profiles and heatmaps were generated using deepTools computeMatrix (scale-regions and reference-point modes; ±1 kb flanking regions; 1 kb scaled gene body; 10 bp bins; --missingDataAsZero), followed by plotProfile and plotHeatmap.

### Small RNA-sequencing and analysis

Sample preparation: Approximately 1,000 Animals were grown on RNAi plates from L4 stages and collected into 1.5mL low-bind Eppendorf’s at the desired age. Animals washed five times in M9 buffer and then snap-frozen in 500uL of Trizol reagent. RNA was extracted as described for RNA-sequencing (see above), where DNAse-treated RNA was then subject to 5’ Pyrophosphohydrolase (RppH; NEB) treatment at 37°C for 30-minutes to remove pyrophosphate from the 5’ end of triphosphorylated RNA. Samples were immediately subject to RNA clean up using isopropanol precipitation overnight at -20°C. Library preparation and sequencing was performed by NovoGene, Oxford, UK.

Analysis: Fastq files were clipped to remove adaptors using fastx_clipper from the Fastx Toolkit (http://hannonlab.cshl.edu/fastx_toolkit/) and converted to fasta files using fastq_to_fasta also from the Fastx Toolkit. Reads were then aligned to the *C. elegans* genome using bowtie with the following parameters: bowtie –f –v 0 –k 1 –S and resultant sam files were converted to bam files using samtools version 1.9 and to bed files using bedtools. Reads were annotated to specific loci using the annotation provided by TinyRNA [108], using bedtools intersect and scripts in R were used to classify reads as microRNAs or 22G-RNAs and to quantify the number of reads. Reads were annotated to specific loci using the annotation provided by TinyRNA [108], using bedtools intersect and scripts in R were used to classify reads as microRNAs or 22G-RNAs and to quantify the number of reads. DESeq2 was used for normalization of the 22G-RNAs and the miRNAs were normalized relative to total read count.

### Outron read mapping

Outron analysis was performed using a custom Python pipeline integrating pyBigWig, NumPy, pandas and SciPy. Gene models were derived from the WBcel235 GTF annotation, and one representative transcript per gene was selected based on maximal exonic length. Analyses were restricted to protein-coding genes with exon length ≥500bp, and genes with TSS overlap (±50bp) or upstream genes within 300bp were excluded. Strand-specific bigWig files (uniquely mapped reads) were used to quantify mean coverage in a defined upstream outron window (150 bp upstream of the TSS) and across the annotated gene body. Reported outron lengths in *C. elegans* can range from 10 to 3,000 nucleotides [109], thus, this window is sufficient to capture any reads mapping to un-processed (i.e., non-trans-spliced) outron regions for analysis.

Outron usage was calculated as the ratio of upstream to gene body density implementing a pseudocount of 0.01. Differential outron usage between conditions was computed as ≥0.58 log2 fold-change of these ratios across biological replicates. Genes were retained if they met minimum coverage thresholds (gene body density ≥0.1, outron density ≥0.01, and ≥5 reads in the outron region in at least one condition). Upstream readthrough was assessed by examining coverage in sequential upstream windows and calculating a regression slope–based readthrough score; genes predicted to exhibit readthrough were excluded from high-confidence analyses.

### Proteomic sample preparation and analysis

On days 4 and 10 of adulthood, 60 animals were collected per condition into 30µL of M9 buffer in protein low-bind tubes. Samples were washed 5-times in 1mL of M9 buffer with brief centrifugation at 500g to pellet worms between washes. Final supernatants were removed, and samples were flash frozen in liquid nitrogen in minimal M9 volume and stored at -70 until further processing. For lysis, 40µL of 7M guanidine hydrochloride in 50mM of ammonia bicarbonate buffer was added, and an equal volume of extraction beads were added to each sample. Sonication was performed in a Biorupter Pico at 4°C using 10-cycles of 30-seconds on/ 30-seconds off, and then centrifuged at 20,000g and 4°C for 15-minutes. Equal amounts of protein (30µg) were then reduced with 5mM Dithiothreitol at 37°C for 1-hour, and immediately alkylated with 20mM iodoacetamide for 30-minutes at 20°C. Trypsin digestion was performed overnight at 37°C with shaking at 650RPM using trypsin: sample ratios of 1:20µg. De-salting was performed using Oasis HLB columns and peptides were quality checked prior to LC-MS/MS analysis.

### LC-MS/MS measurement

Peptides were separated by nano-liquid chromatography (Thermo Fisher Scientific Ultimate RSLC 3000) coupled in line a Q Exactive mass spectrometer equipped with an Easy-Spray source (Thermo Fischer Scientific). Peptides were trapped onto a C18 PepMac100 precolumn (300µm i.d.x 5mm, 100Å, ThermoFischer Scientific) using Solvent A (0.1% Formic acid, HPLC grade water). The peptides were further separated onto an Easy-Spray RSLC C18 column (75um i.d., 50cm length, Thermo Fischer Scientific) using a 60 minutes linear gradient (15% to 35% solvent B (0.1% formic acid in acetonitrile)) at a flow rate 200nl/min. The raw data were acquired on the mass spectrometer in a data-dependent acquisition mode (DDA). Full-scan MS spectra were acquired in the Orbitrap (Scan range 350-1500m/z, resolution 70,000; AGC target, 3e6, maximum injection time, 50ms). The 10 most intense peaks were selected for higher-energy collision dissociation (HCD) fragmentation at 30% of normalized collision energy. HCD spectra were acquired in the Orbitrap at resolution 17,500, AGC target 5e4, maximum injection time 120ms with fixed mass at 180m/z. Charge exclusion was selected for unassigned and 1+ ions. The dynamic exclusion was set to 20s.

### Statistical analyses

All gene ontology enrichment analyses were performed by searching gene sets within the g:Profiler platform [110] with significant terms defined if adjusted P-values ≤0.05. Figures and statistics were curated and analysed using Graphpad Prism (version 10), Rstudio and deepTools. Final figures were curated in Adobe illustrator (2025). Schematics were made using Biorender.

## Supplementary figures and legends

**Supplemental figure 1.**
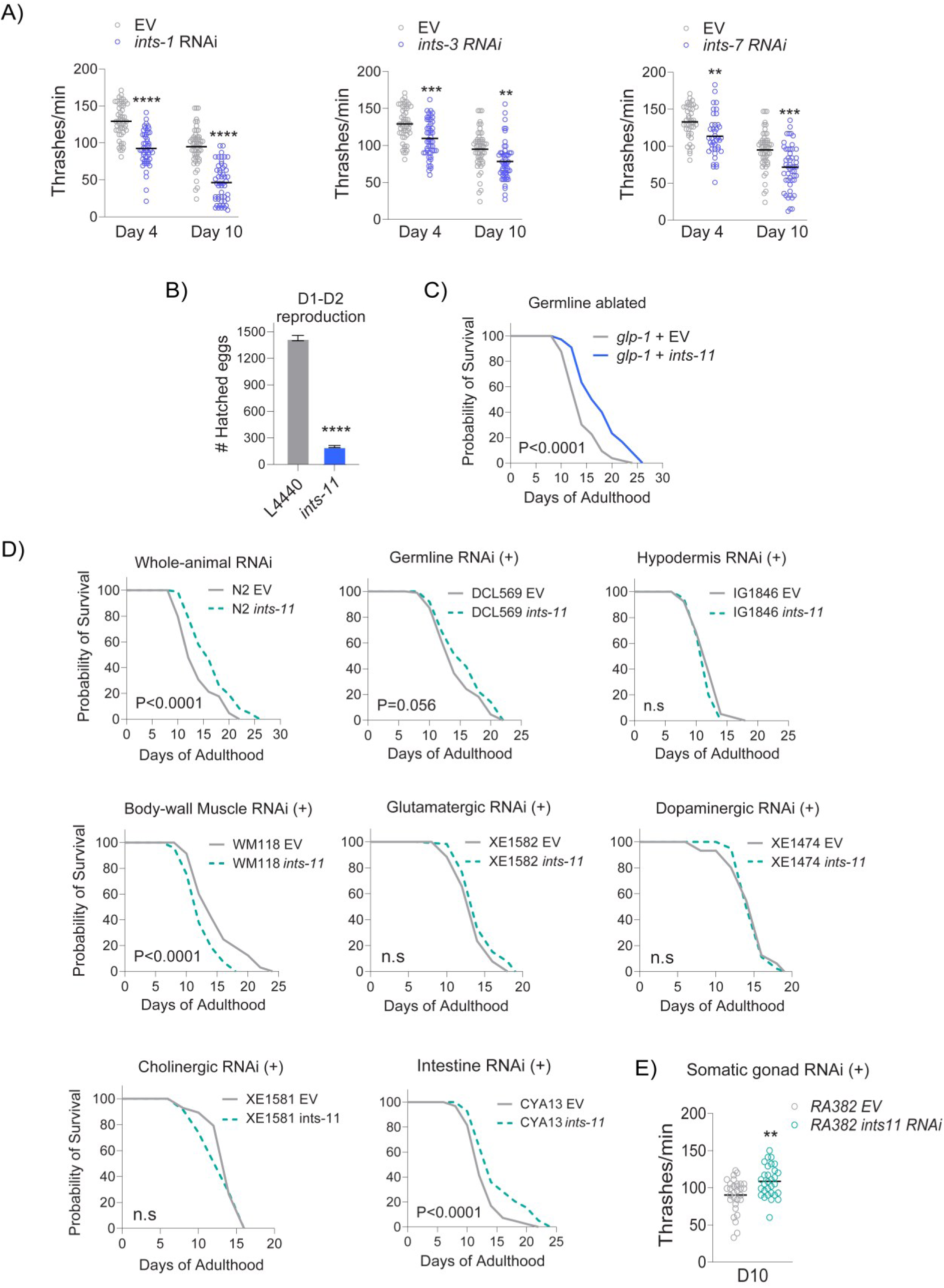
**A)** Thrashing performance in day-4 and day-10 animals showing impaired healthspan from adulthood depletion of *ints-1, ints-3* and *ints-7* Integrator subunits. **B)** Egg hatching assay from 5 combined representative individuals fed either empty vector or *ints-11* dsRNA from L4 stage. **C)** RNAi against *ints-11* within sterile *glp-1(e2141)* mutants extends lifespan. **D)** Tissue-specific RNAi screen for *ints-11*. **E)** Thrashing rates are enhanced from *ints-11* RNAi within the somatic gonad. Thrashing assays are from 2 independent repeats containing ∼50 individuals. Lifespans are biological repeats from ∼200 individuals.

**Supplemental figure 2.**
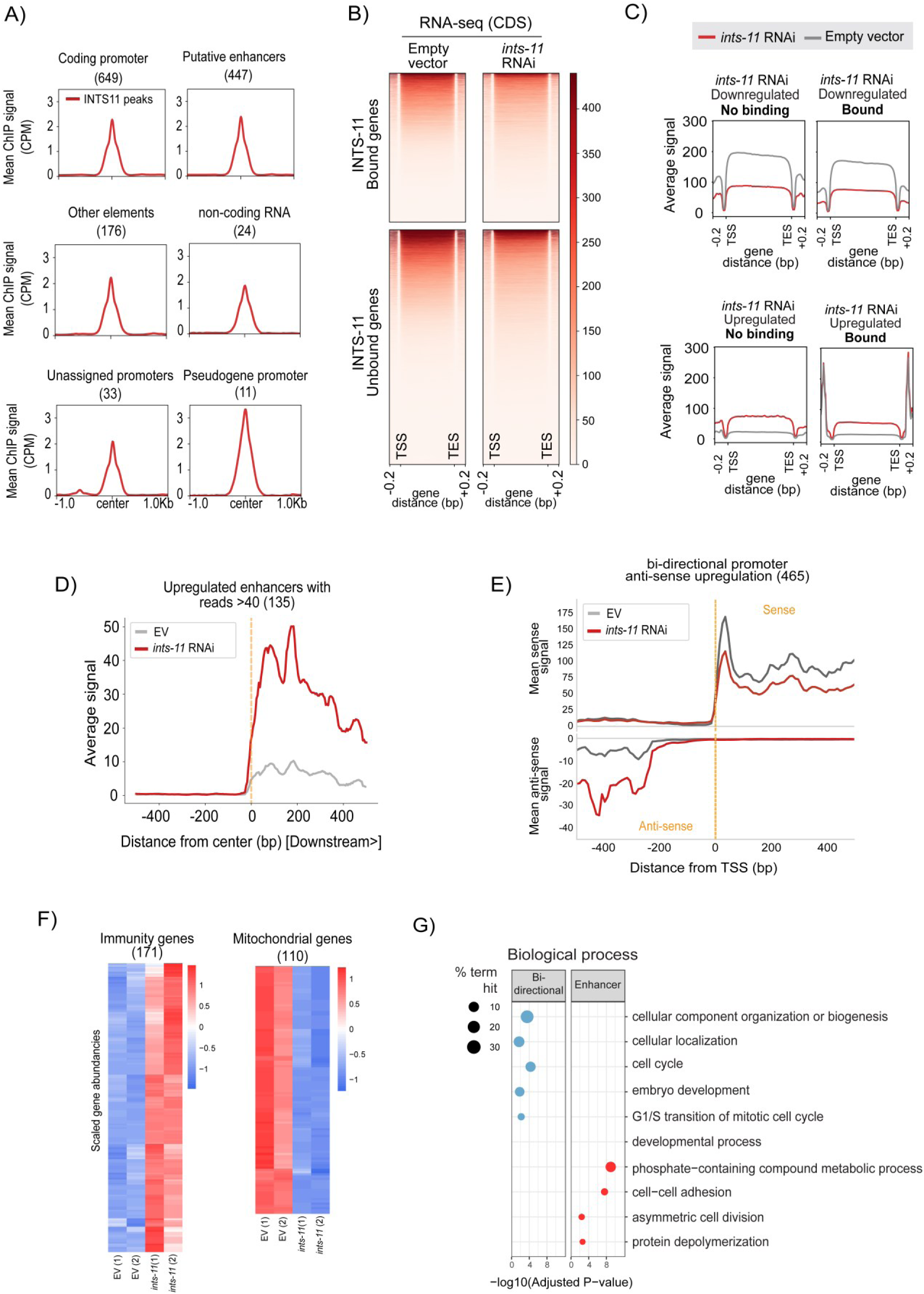
**A)** Mean chIP-centred peak intensity plots across multiple regulatory transcriptional loci. **B)** Heatmap comparing RNA-seq expression coverage across coding domains of INTS-11 bound versus unbound genes in empty vector (left) and *ints-11* RNAi-fed (right) animals. **C)** Metagene plots across coding domains of INTS11 bound versus unbound genes, but stratified into subsets identified as either differentially up/downregulated within RNA-seq data. **D)** Metagene plot for upregulated enhancer loci transcription for genes with moderate (>40 average reads) RNA-seq expression coverage. **E)** RNA-seq metagene plot for bidirectional promoter antisense-upregulated protein-coding genes. **F)** Scaled heatmaps of mitochondria- and immune-related genes from differentially expressed RNA-seq data. **G)** Gene ontology enrichment of genes associated with upregulated enhancer (left panel) and bidirectional promoter (right panel).

**Supplemental figure 3.**
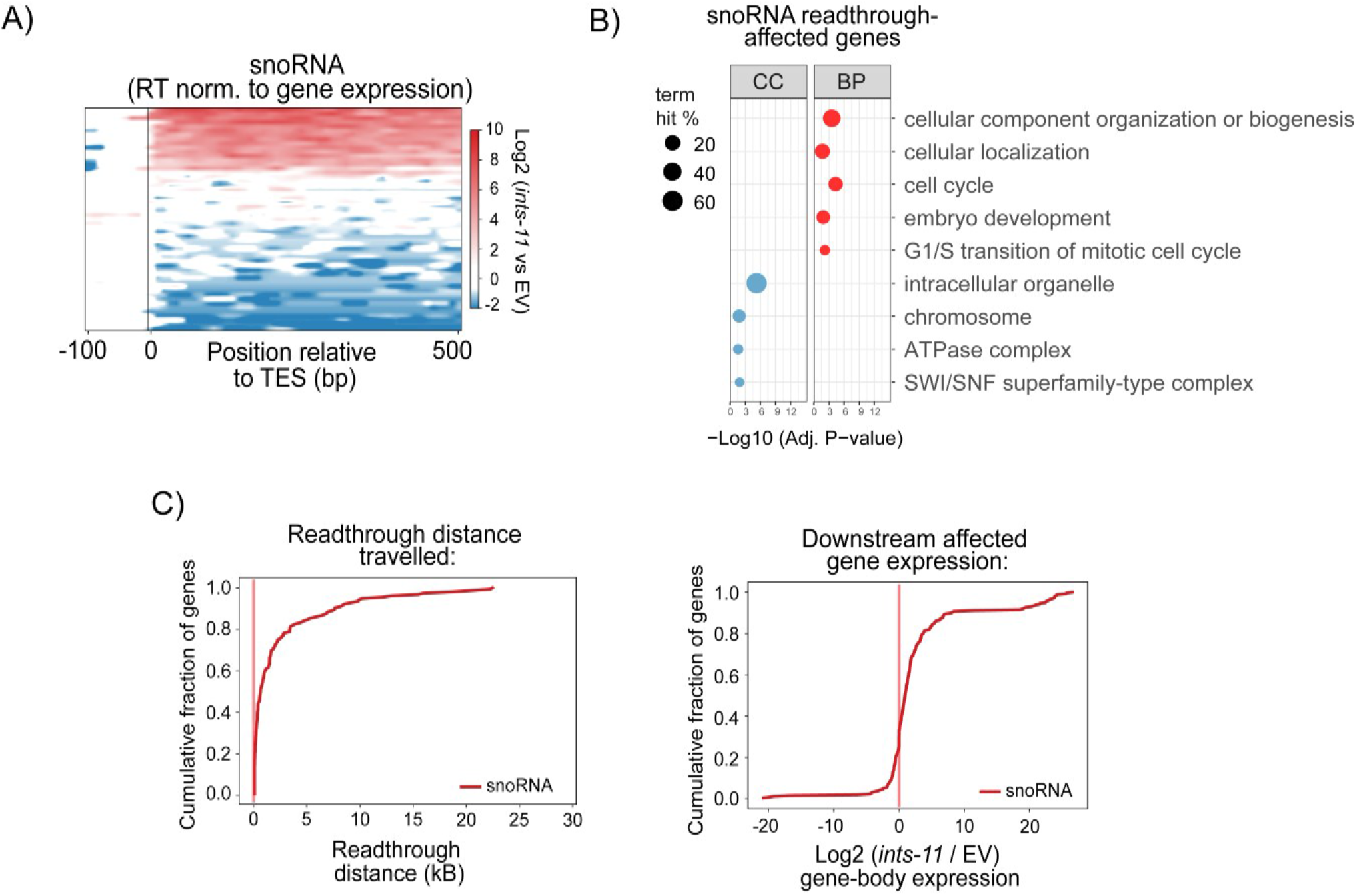
**A)** Heatmap of pol II readthrough events at snoRNA TES loci. **B)** Gene ontology enrichment analysis of snoRNA readthrough-affected genes. **C)** Cumulative distribution plot of distance travelled from snoRNA readthrough genes (left), and gene expression of downstream affected genes residing within snoRNA readthrough windows (right).

**Supplemental figure 4.**
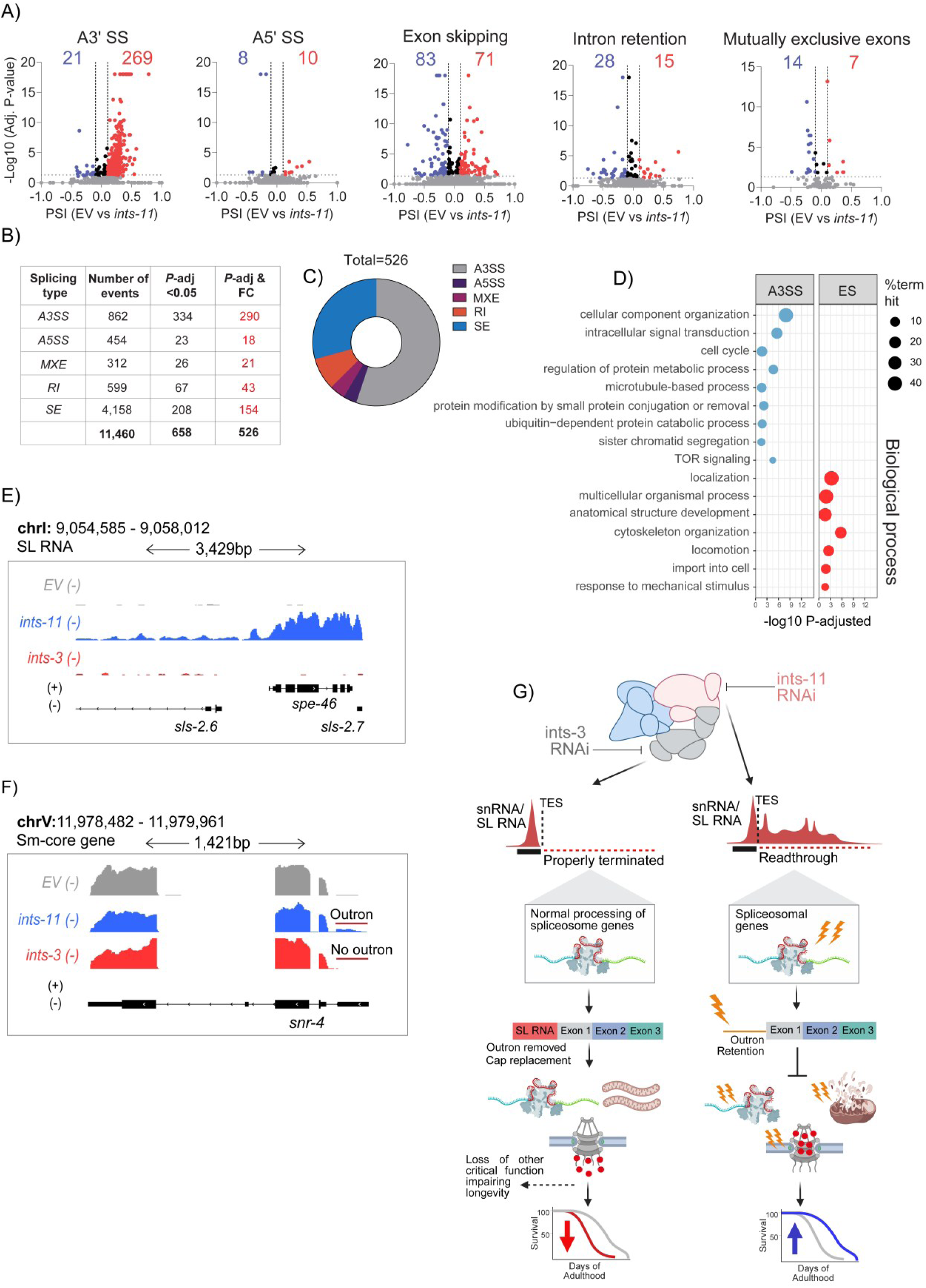
**A)** Volcano plots representing percentage spliced in (PSI) for alternatively spliced isoforms identified within RNA-seq datasets. **B)** Summary table for differential splicing events. **C)** Pie chart summarising distribution of alternative splicing events across splicing classes. **D)** Gene ontology enrichment for alternatively spliced events. **E)** Genome coverage tracks highlighting lack of SL RNA termination defects from *ints-3* RNAi (negatively effects lifespan), but stark readthrough from *ints-11* RNAi (lifespan extending effects). **F)** Genome coverage tracks highlighting the presence/ absence of outron retention from *ints-11* versus *ints-3* RNAi, respectively. **G)** Model indicating defective termination of sn/SL RNA and consequential impacts on spliceosomal genes sit upstream of other contributing longevity programmes from loss of *ints-11*.

**Supplemental figure 5.**
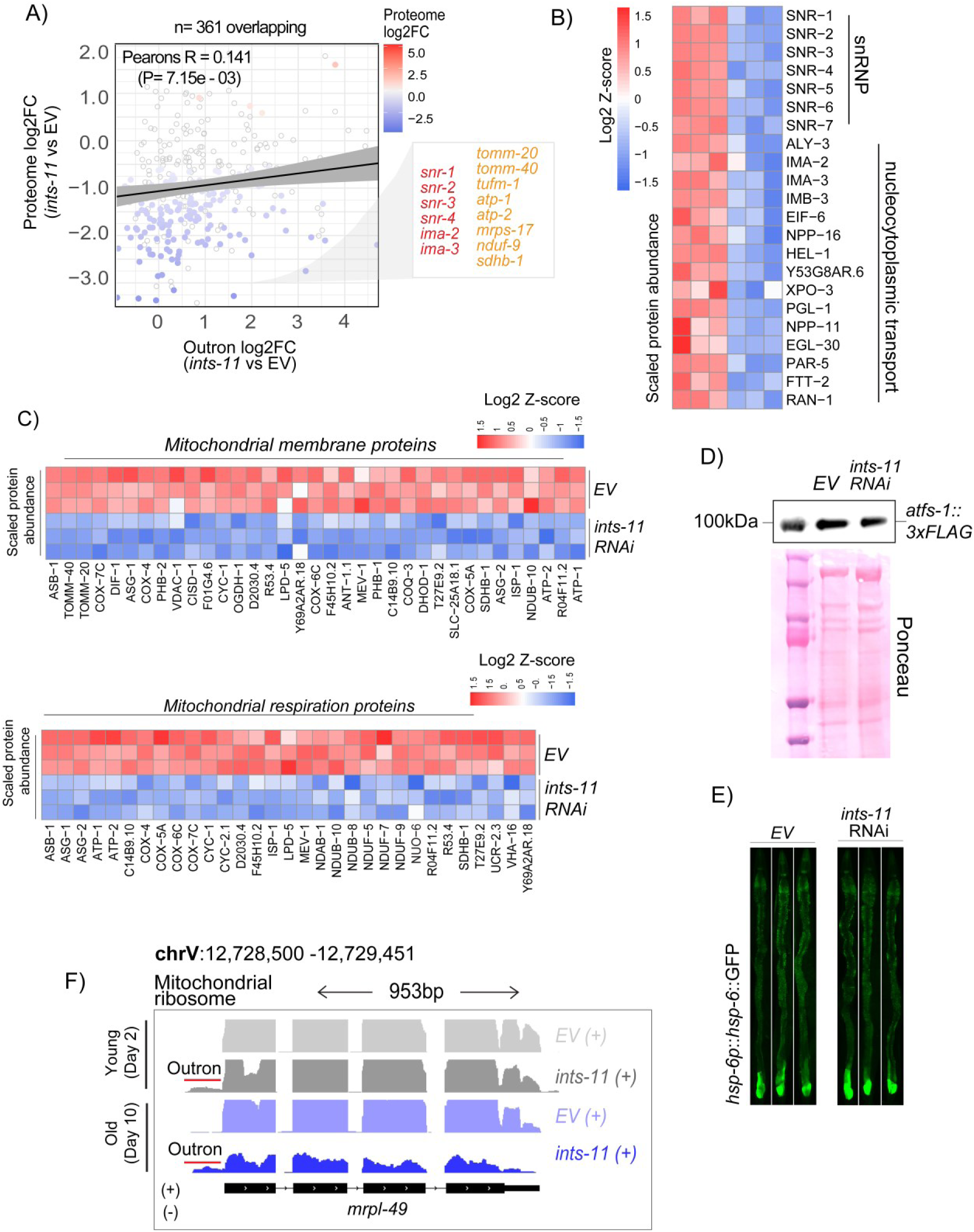
**A)** Correlation plot comparing outron-retaining genes with protein abundance levels from whole-animal proteomics. **B)** Log2 Z-score scaled heatmaps of core spliceosomal and nucleocytoplasmic proteins implicated in trans-splicing and snRNP biogenesis pathways. **C)** Heatmaps of mitochondrial outer-membrane (top) and respiratory (bottom) proteins. **D)** Western blot of an endogenously-tagged ATFS-1 strain after 5-days of *ints-11* RNAi, with ponceau staining as a representative loading control. **E)** Representative microscopy images of the *phsp-6::hsp-6::GFP* reporter strain after 5-days of *ints-11* RNAi. **F)** Genome browser tracks of a mitochondrial ribosome gene exhibiting outron retention in day-2 and day-10 *ints-11* silenced animals.

## Notes

### Competing Interest Statement

The authors have declared no competing interest.

## References

1. Our Ageing Population, The State of Ageing 2025. Available from: https://ageing-better.org.uk/.

2. Frenk, S. and J. Houseley, Gene expression hallmarks of cellular ageing. Biogerontology, 2018. 19(6): p. 547–566.

3. Harries, L.W., Dysregulated RNA processing and metabolism: a new hallmark of ageing and provocation for cellular senescence. FEBS J, 2023. 290(5): p. 1221–1234.

4. Osman, S. and P. Cramer, Structural Biology of RNA Polymerase II Transcription: 20 Years On. Annu Rev Cell Dev Biol, 2020. 36: p. 1–34.

5. Kus, K., et al., DSIF factor Spt5 coordinates transcription, maturation and exoribonucleolysis of RNA polymerase II transcripts. Nat Commun, 2025. 16(1): p. 10.

6. Debes, C., et al., Ageing-associated changes in transcriptional elongation influence longevity. Nature, 2023. 616(7958): p. 814–821.

7. Rangaraju, S., et al., Correction: Suppression of transcriptional drift extends C. elegans lifespan by postponing the onset of mortality. Elife, 2016. 5.

8. Tatomer, D.C., et al., The Integrator complex cleaves nascent mRNAs to attenuate transcription. Genes Dev, 2019. 33(21-22): p. 1525–1538.

9. Lai, F., et al., Integrator mediates the biogenesis of enhancer RNAs. Nature, 2015. 525(7569): p. 399–403.

10. Gardini, A., et al., Integrator regulates transcriptional initiation and pause release following activation. Mol Cell, 2014. 56(1): p. 128–139.

11. Yue, J., et al., Integrator orchestrates RAS/ERK1/2 signaling transcriptional programs. Genes Dev, 2017. 31(17): p. 1809–1820.

12. Barbieri, E., et al., Targeted Enhancer Activation by a Subunit of the Integrator Complex. Mol Cell, 2018. 71(1): p. 103–116 e7.

13. Otani, Y., et al., Integrator complex plays an essential role in adipose differentiation. Biochem Biophys Res Commun, 2013. 434(2): p. 197–202.

14. Confino, S., et al., INTS1 is required for maintaining accurate transcriptional integrity and behavior in zebrafish. NAR Genom Bioinform, 2025. 7(4): p. lqaf185.

15. Baluapuri, A., et al., Integrator loss leads to dsRNA formation that triggers the integrated stress response. Cell, 2025. 188(12): p. 3184–3201 e21.

16. Baillat, D., et al., Integrator, a multiprotein mediator of small nuclear RNA processing, associates with the C-terminal repeat of RNA polymerase II. Cell, 2005. 123(2): p. 265–76.

17. Dasilva, L.F., et al., Integrator enforces the fidelity of transcriptional termination at protein-coding genes. Sci Adv, 2021. 7(45): p. eabe3393.

18. Elrod, N.D., et al., The Integrator Complex Attenuates Promoter-Proximal Transcription at Protein-Coding Genes. Mol Cell, 2019. 76(5): p. 738–752 e7.

19. Gomez-Orte, E., et al., Disruption of the Caenorhabditis elegans Integrator complex triggers a non-conventional transcriptional mechanism beyond snRNA genes. PLoS Genet, 2019. 15(2): p. e1007981.

20. Eaton, J.D., et al., Human promoter directionality is determined by transcriptional initiation and the opposing activities of INTS11 and CDK9. Elife, 2024. 13.

21. Yang, J., et al., Transcription directionality is licensed by Integrator at active human promoters. Nat Struct Mol Biol, 2024. 31(8): p. 1208–1221.

22. Vervoort, S.J., et al., The PP2A-Integrator-CDK9 axis fine-tunes transcription and can be targeted therapeutically in cancer. Cell, 2021. 184(12): p. 3143–3162 e32.

23. Huang, K.L., et al., Integrator Recruits Protein Phosphatase 2A to Prevent Pause Release and Facilitate Transcription Termination. Mol Cell, 2020. 80(2): p. 345–358 e9.

24. Zheng, H., et al., Identification of Integrator-PP2A complex (INTAC), an RNA polymerase II phosphatase. Science, 2020. 370(6520).

25. Fianu, I., et al., Structural basis of Integrator-dependent RNA polymerase II termination. Nature, 2024. 629(8010): p. 219–227.

26. Rubtsova, M.P., et al., Integrator is a key component of human telomerase RNA biogenesis. Sci Rep, 2019. 9(1): p. 1701.

27. Beckedorff, F., et al., The Human Integrator Complex Facilitates Transcriptional Elongation by Endonucleolytic Cleavage of Nascent Transcripts. Cell Rep, 2020. 32(3): p. 107917.

28. Gurumurthy, A., et al., Super-enhancer mediated regulation of adult beta-globin gene expression: the role of eRNA and Integrator. Nucleic Acids Res, 2021. 49(3): p. 1383–1396.

29. Xie, M., et al., The host Integrator complex acts in transcription-independent maturation of herpesvirus microRNA 3’ ends. Genes Dev, 2015. 29(14): p. 1552–64.

30. Kirstein, N., et al., The Integrator complex regulates microRNA abundance through RISC loading. Sci Adv, 2023. 9(6): p. eadf0597.

31. Cazalla, D., M. Xie, and J.A. Steitz, A primate herpesvirus uses the integrator complex to generate viral microRNAs. Mol Cell, 2011. 43(6): p. 982–92.

32. Beltran, T., et al., Integrator is recruited to promoter-proximally paused RNA Pol II to generate Caenorhabditis elegans piRNA precursors. EMBO J, 2021. 40(5): p. e105564.

33. Czech, B., et al., A transcriptome-wide RNAi screen in the Drosophila ovary reveals factors of the germline piRNA pathway. Mol Cell, 2013. 50(5): p. 749–61.

34. Fianu, I., et al., Structural basis of Integrator-mediated transcription regulation. Science, 2021. 374(6569): p. 883–887.

35. Pfleiderer, M.M. and W.P. Galej, Structure of the catalytic core of the Integrator complex. Mol Cell, 2021. 81(6): p. 1246–1259 e8.

36. Razew, M., et al., Structural basis of the Integrator complex assembly and association with transcription factors. Mol Cell, 2024. 84(13): p. 2542–2552 e5.

37. Waddell, B.M., et al., Differential effect of ubiquitous and germline depletion of Integrator complex function on C. elegans physiology. Biol Open, 2025. 14(4).

38. Waddell, B.M. and C.W. Wu, A role for the C. elegans Argonaute protein CSR-1 in small nuclear RNA 3’ processing. PLoS Genet, 2024. 20(5): p. e1011284.

39. Cihlarova, Z., et al., BRAT1 links Integrator and defective RNA processing with neurodegeneration. Nat Commun, 2022. 13(1): p. 5026.

40. Lim, B., et al., Genetic alterations and their clinical implications in gastric cancer peritoneal carcinomatosis revealed by whole-exome sequencing of malignant ascites. Oncotarget, 2016. 7(7): p. 8055–66.

41. Simpson, H.M., et al., Concurrent Mutations in ATM and Genes Associated with Common gamma Chain Signaling in Peripheral T Cell Lymphoma. PLoS One, 2015. 10(11): p. e0141906.

42. Li, X., et al., The pro-tumor activity of INTS7 on lung adenocarcinoma via inhibiting immune infiltration and activating p38MAPK pathway. Sci Rep, 2024. 14(1): p. 25636.

43. Tayari, M.M. and R. Shiekhattar, Integrator 20th anniversary: a molecular machine indispensable in development and disease. Trends Mol Med, 2025.

44. Wang, B., et al., Comprehensive analysis of INTS family related to expression, prognosis, diagnosis and immune features in hepatocellular carcinoma. Heliyon, 2024. 10(9): p. e30244.

45. Kenyon, C., et al., A C. elegans mutant that lives twice as long as wild type. Nature, 1993. 366(6454): p. 461–4.

46. Lin, K., et al., daf-16: An HNF-3/forkhead family member that can function to double the life-span of Caenorhabditis elegans. Science, 1997. 278(5341): p. 1319–22.

47. Vellai, T., et al., Genetics: influence of TOR kinase on lifespan in C. elegans. Nature, 2003. 426(6967): p. 620.

48. Berkyurek, A.C., et al., The RNA polymerase II subunit RPB-9 recruits the integrator complex to terminate Caenorhabditis elegans piRNA transcription. EMBO J, 2021. 40(5): p. e105565.

49. de la Guardia, Y., et al., Run-on of germline apoptosis promotes gonad senescence in C. elegans. Oncotarget, 2016. 7(26): p. 39082–39096.

50. Ezcurra, M., et al., C. elegans Eats Its Own Intestine to Make Yolk Leading to Multiple Senescent Pathologies. Curr Biol, 2018. 28(20): p. 3352.

51. Gems, D. and Y. de la Guardia, Alternative Perspectives on Aging in Caenorhabditis elegans: Reactive Oxygen Species or Hyperfunction? Antioxid Redox Signal, 2013. 19(3): p. 321–9.

52. Slade, L., T. Etheridge, and N.J. Szewczyk, Consolidating multiple evolutionary theories of ageing suggests a need for new approaches to study genetic contributions to ageing decline. Ageing Res Rev, 2024. 100: p. 102456.

53. Wang, H., et al., A parthenogenetic quasi-program causes teratoma-like tumors during aging in wild-type C. elegans. NPJ Aging Mech Dis, 2018. 4: p. 6.

54. Williams, G.C., Pleiotropy, Natural Selection, and the Evolution of Senescence. Evolution, 1957. 11(4): p. 398–411.

55. Kirkwood, T.B., Evolution of ageing. Nature, 1977. 270(5635): p. 301–4.

56. Rose, M. and B. Charlesworth, A test of evolutionary theories of senescence. Nature, 1980. 287(5778): p. 141–2.

57. Rose, M.R. and B. Charlesworth, Genetics of life history in Drosophila melanogaster. II. Exploratory selection experiments. Genetics, 1981. 97(1): p. 187–96.

58. Wattiaux, J.M., Parental age effects in Drosophila pseudoobscura. Exp Gerontol, 1968. 3(1): p. 55–61.

59. Janes, J., et al., Chromatin accessibility dynamics across C. elegans development and ageing. Elife, 2018. 7.

60. Sabath, K., et al., Basis of gene-specific transcription regulation by the Integrator complex. Mol Cell, 2024. 84(13): p. 2525–2541 e12.

61. Stadelmayer, B., et al., Integrator complex regulates NELF-mediated RNA polymerase II pause/release and processivity at coding genes. Nat Commun, 2014. 5: p. 5531.

62. Narita, T., et al., Human transcription elongation factor NELF: identification of novel subunits and reconstitution of the functionally active complex. Mol Cell Biol, 2003. 23(6): p. 1863–73.

63. Kowalczyk, M.S., et al., Intragenic enhancers act as alternative promoters. Mol Cell, 2012. 45(4): p. 447–58.

64. Cinghu, S., et al., Intragenic Enhancers Attenuate Host Gene Expression. Mol Cell, 2017. 68(1): p. 104–117 e6.

65. Chen, R.A., et al., The landscape of RNA polymerase II transcription initiation in C. elegans reveals promoter and enhancer architectures. Genome Res, 2013. 23(8): p. 1339–47.

66. Ntini, E., et al., Polyadenylation site-induced decay of upstream transcripts enforces promoter directionality. Nat Struct Mol Biol, 2013. 20(8): p. 923–8.

67. Preker, P., et al., RNA exosome depletion reveals transcription upstream of active human promoters. Science, 2008. 322(5909): p. 1851-4.

68. Hobson, D.J., et al., RNA polymerase II collision interrupts convergent transcription. Mol Cell, 2012. 48(3): p. 365–74.

69. Shearwin, K.E., B.P. Callen, and J.B. Egan, Transcriptional interference--a crash course. Trends Genet, 2005. 21(6): p. 339–45.

70. Blumenthal, T., Trans-splicing and operons in C. elegans. WormBook, 2012: p. 1–11.

71. Graber, J.H., et al., C. elegans sequences that control trans-splicing and operon pre-mRNA processing. RNA, 2007. 13(9): p. 1409–26.

72. Wang, Y., et al., rMATS-turbo: an efficient and flexible computational tool for alternative splicing analysis of large-scale RNA-seq data. Nat Protoc, 2024. 19(4): p. 1083–1104.

73. Cheng, G., et al., In vivo translation and stability of trans-spliced mRNAs in nematode embryos. Mol Biochem Parasitol, 2007. 153(2): p. 95–106.

74. MacMorris, M., et al., A novel family of C. elegans snRNPs contains proteins associated with trans-splicing. RNA, 2007. 13(4): p. 511–20.

75. Coy, S., et al., The Sm complex is required for the processing of non-coding RNAs by the exosome. PLoS One, 2013. 8(6): p. e65606.

76. Ma, T., et al., Sm complex assembly and 5’ cap trimethylation promote selective processing of snRNAs by the 3’ exonuclease TOE1. Proc Natl Acad Sci U S A, 2024. 121(3): p. e2315259121.

77. Seipelt, R.L., et al., U1 snRNA is cleaved by RNase III and processed through an Sm site-dependent pathway. Nucleic Acids Res, 1999. 27(2): p. 587–95.

78. Philippe, L., et al., An in vivo genetic screen for genes involved in spliced leader trans-splicing indicates a crucial role for continuous de novo spliced leader RNP assembly. Nucleic Acids Res, 2017. 45(14): p. 8474–8483.

79. Yang, Y.F., et al., Trans-splicing enhances translational efficiency in C. elegans. Genome Res, 2017. 27(9): p. 1525–1535.

80. Durieux, J., S. Wolff, and A. Dillin, The cell-non-autonomous nature of electron transport chain-mediated longevity. Cell, 2011. 144(1): p. 79–91.

81. Charmpilas, N., et al., Reproductive regulation of the mitochondrial stress response in Caenorhabditis elegans. Cell Rep, 2024. 43(6): p. 114336.

82. Diodato, D., D. Ghezzi, and V. Tiranti, The Mitochondrial Aminoacyl tRNA Synthetases: Genes and Syndromes. Int J Cell Biol, 2014. 2014: p. 787956.

83. Zheng, T., et al., Cytoplasmic and mitochondrial aminoacyl-tRNA synthetases differentially regulate lifespan in Caenorhabditis elegans. iScience, 2022. 25(11): p. 105266.

84. Hu, I.M., et al., Immuno-metabolic stress responses control longevity from mitochondrial translation inhibition in C. elegans. Nat Commun, 2025. 16(1): p. 6083.

85. Yang, W. and S. Hekimi, Two modes of mitochondrial dysfunction lead independently to lifespan extension in Caenorhabditis elegans. Aging Cell, 2010. 9(3): p. 433–47.

86. de Lencastre, A., et al., MicroRNAs both promote and antagonize longevity in C. elegans. Curr Biol, 2010. 20(24): p. 2159–68.

87. Pincus, Z., T. Smith-Vikos, and F.J. Slack, MicroRNA predictors of longevity in Caenorhabditis elegans. PLoS Genet, 2011. 7(9): p. e1002306.

88. Dexheimer, P.J., J. Wang, and L. Cochella, Two MicroRNAs Are Sufficient for Embryonic Patterning in C. elegans. Curr Biol, 2020. 30(24): p. 5058–5065 e5.

89. Lehrbach, N.J., et al., Post-developmental microRNA expression is required for normal physiology, and regulates aging in parallel to insulin/IGF-1 signaling in C. elegans. RNA, 2012. 18(12): p. 2220–35.

90. Gu, W., et al., Distinct argonaute-mediated 22G-RNA pathways direct genome surveillance in the C. elegans germline. Mol Cell, 2009. 36(2): p. 231–44.

91. Mao, K., P. Breen, and G. Ruvkun, Mitochondrial dysfunction induces RNA interference in C. elegans through a pathway homologous to the mammalian RIG-I antiviral response. PLoS Biol, 2020. 18(12): p. e3000996.

92. Zhang, C., et al., mut-16 and other mutator class genes modulate 22G and 26G siRNA pathways in Caenorhabditis elegans. Proc Natl Acad Sci U S A, 2011. 108(4): p. 1201–8.

93. Maxwell, C.S., et al., Pol II docking and pausing at growth and stress genes in C. elegans. Cell Rep, 2014. 6(3): p. 455–66.

94. Yamamoto, J., et al., DSIF and NELF interact with Integrator to specify the correct post-transcriptional fate of snRNA genes. Nat Commun, 2014. 5: p. 4263.

95. Dillin, A., et al., Rates of behavior and aging specified by mitochondrial function during development. Science, 2002. 298(5602): p. 2398–401.

96. Jensen, M.B. and H. Jasper, Mitochondrial proteostasis in the control of aging and longevity. Cell Metab, 2014. 20(2): p. 214–25.

97. Takata, H., et al., The integrator complex is required for integrity of Cajal bodies. J Cell Sci, 2012. 125(Pt 1): p. 166–75.

98. Liu, Y., A. Beyer, and R. Aebersold, On the Dependency of Cellular Protein Levels on mRNA Abundance. Cell, 2016. 165(3): p. 535–50.

99. Vogel, C. and E.M. Marcotte, Insights into the regulation of protein abundance from proteomic and transcriptomic analyses. Nat Rev Genet, 2012. 13(4): p. 227–32.

100. Skaar, J.R., et al., INTS3 controls the hSSB1-mediated DNA damage response. J Cell Biol, 2009. 187(1): p. 25–32.

101. Stiernagle, T., Maintenance of C. elegans. WormBook, 2006: p. 1–11.

102. Qadota, H., et al., Establishment of a tissue-specific RNAi system in C. elegans. Gene, 2007. 400(1-2): p. 166–73.

103. Watts, J.S., et al., New Strains for Tissue-Specific RNAi Studies in Caenorhabditis elegans. G3 (Bethesda), 2020. 10(11): p. 4167–4176.

104. Smith, H.J., et al., Neuronal mTORC1 inhibition promotes longevity without suppressing anabolic growth and reproduction in C. elegans. PLoS Genet, 2023. 19(9): p. e1010938.

105. Fischer, C.A., et al., MitoSegNet: Easy-to-use Deep Learning Segmentation for Analyzing Mitochondrial Morphology. iScience, 2020. 23(10): p. 101601.

106. Love, M.I., W. Huber, and S. Anders, Moderated estimation of fold change and dispersion for RNA-seq data with DESeq2. Genome Biol, 2014. 15(12): p. 550.

107. Sen, I., A. Kavsek, and C.G. Riedel, Chromatin Immunoprecipitation and Sequencing (ChIP-seq) Optimized for Application in Caenorhabditis elegans. Curr Protoc, 2021. 1(7): p. e187.

108. Tate, A.J., K.C. Brown, and T.A. Montgomery, tiny-count: a counting tool for hierarchical classification and quantification of small RNA-seq reads with single-nucleotide precision. Bioinform Adv, 2023. 3(1): p. vbad065.

109. Saito, T.L., et al., The transcription start site landscape of C. elegans. Genome Res, 2013. 23(8): p. 1348–61.

110. Raudvere, U., et al., g:Profiler: a web server for functional enrichment analysis and conversions of gene lists (2019 update). Nucleic Acids Res, 2019. 47(W1): p. W191–W198.

